# Learning regularities in noise engages both neural predictive activity and representational changes

**DOI:** 10.1101/2025.08.18.670891

**Authors:** Coumarane Tirou, Oussama Abdoun, Teodóra Vékony, Laure Tosatto, Andrea Brovelli, Marine Vernet, Dezső Németh, Romain Quentin

**Affiliations:** Université Claude Bernard Lyon 1, INSERM U1028, CNRS UMR5292, Lyon Neuroscience Research Center (CRNL), EDUWELL team, Lyon, France; Gran Canaria Cognitive Research Center, Atlántico Medio University, Las Palmas de Gran Canaria, Spain; School of Biological Sciences, Monash University, Melbourne, VIC 3800, Australia; Normandie University, UNICAEN, CNRS, EthoS, UMR 6552, 14000 Caen, France; Institut de Neurosciences de la Timone UMR 7289, Aix Marseille Université, CNRS, 13005, Marseille, France; Université Claude Bernard Lyon 1, INSERM U1028, CNRS UMR5292, Lyon Neuroscience Research Center (CRNL), IMPACT team, Lyon, France; BML-NAP Research Group, ELTE Eötvös Loránd University & HUN-REN Research Centre for Natural Sciences, Budapest, Hungary

**Author notes:** These authors jointly supervised this work (Dezső Németh, Romain Quentin).

## Abstract

The ability to extract structured sensory patterns from a noisy environment is fundamental to cognition, yet how the brain learns complex regularities remains unclear. Using magnetoencephalography during a visuomotor task, we tracked the neural dynamics as humans learned non-adjacent temporal dependencies embedded in noise. We reveal that learning is supported by two temporally dissociable mechanisms. Neural predictive activity emerged rapidly, with stimulus-specific patterns appearing before stimulus onset and preceding measurable behavioral improvements. This is followed by a slower build-up of representational change, characterized by an increased neural pattern similarity between statistically dependent, non-adjacent elements. Both processes are supported by a distributed consortium of networks, with the sensorimotor and dorsal attentional networks playing a central role. These findings suggest that both neural predictive activity and representational changes contribute to learning regularities, revealing a temporal hierarchy in which neural predictive activity precedes behavioral improvement and is followed by neural representational changes, possibly facilitating the gradual consolidation of knowledge into stable neural representations.

## Introduction

Learning allows humans to adapt to a constantly changing environment. Statistical learning plays a key role in this process as it enables the extraction of regularities from the environment without feedback or reward^1^. As a fundamental function of humans’ cognition, such learning underlies a myriad of behaviors including language processing, skill acquisition, or perceptual decision making^2^. Statistical learning^3^ has been extensively studied^3–6^ and shown to be a domain-general mechanism operating across multiple modalities^7–11^. It plays a key role in natural language processing^3,11,12^, as the hierarchical nature of language allows words to form relationships over varying distances within a sentence, resulting in non-adjacent dependencies^4,5^. For example, in the sentence *The book that the girl read was interesting*, the verb *was* must be linked to *book* despite the intervening phrase *that the girl read*. Unlike other sequential structures such as music, where adjacent relationships (e.g., between successive notes) often dominate, everyday communication is filled with such long-distance connections. Processing non-adjacent dependencies is therefore a hallmark of human language expertise.

Studies in humans have shown that extracting regularities is more challenging for non-adjacent than for adjacent dependencies^5,7,8,13^ and that sensitivity to non-adjacent dependencies develops later in infancy^7,9^. Existing theories posit that the learning of adjacent dependencies relies primarily on bottom-up signals from perceptual processing networks, whereas the learning of non-adjacent dependencies involves top-down signals from cognitive control networks to inhibit attention to distracting items and redirect it toward the relevant dependencies^7,10^. Recent advancements in neuroimaging and signal processing^7^ have provided mechanistic insights into the neural mechanisms involved in statistical learning of adjacent dependencies, namely predictive activity and representational change^12,14–16^. These putative mechanisms are neither independent nor mutually exclusive. They have been shown to be dissociable^14^ yet to interact synergistically, with predictive processing shaping the representational geometry of statistical regularities^15^. However, it remains unclear whether these mechanisms also support statistical learning of non-adjacent dependencies^11^, and how they dynamically unfold during learning.

One candidate mechanism is predictive processing^17–20^, which posits that brains are predictive machines^17^ that constantly try to match incoming sensory inputs. The Bayesian brain theory of predictive coding postulates that the brain constructs an internal model of the world by detecting statistical regularities from sensory inputs^21^. Under this theory, the brain has priors on the environment based on its internal model that it uses to make predictions (i.e., top-down modulations) about upcoming stimuli. When an unexpected event arises, it makes use of prediction errors to update its internal model (i.e., through bottom-up modulations) thereby continuously improving its predictions. Predictive coding has been extensively studied across sensory modalities with neural recordings^22,23^ in which distinct brain responses are observed to an expected and a deviant stimulus in sensory regions (e.g., visual or auditory). Some studies^24,25^ suggest that the brain pre-activates neural ensembles involved in the perception of an upcoming, predictable stimulus before it appears (e.g., predicting a tone at 440 Hz activates similar patterns of brain activity as perceiving it). However, evidence for this hypothesis remains scarce, and methodological limitations identified by our research group^26^ have compromised such existing evidence^25^. Thus, it remains unclear whether predictive activity emerges and how it builds up over time during the learning of non-adjacent dependencies.

The other candidate mechanism that may explain statistical learning at the brain level is neural representational change, which is defined as the increase in similarity of neural activity patterns between associated stimuli^12,14–16^. A seminal study in monkeys^27^ demonstrated that neurons in the inferior temporal lobe developed similar responses to images presented in close temporal proximity, suggesting that neural selectivity can emerge from learned temporal associations and may reflect a mechanism for statistical learning. A similar effect has been observed in humans using fMRI^14^, showing that passive exposure to structured and random sequences of abstract images led to increased neural pattern similarity for structured versus random image pairs in the medial temporal lobe and hippocampus. Studies examining changes in neural similarity typically focused on post-learning effects, without measuring behavioral evidence of learning (e.g., reaction time or error rates). They did not track neural changes as learning unfolded, nor did they link these changes to behavioral markers of learning^14,28^. Similarly to predictive activity, it remains unclear whether representational change emerges during the learning of non-adjacent dependencies, and if so, how it develops over time.

Using an explicit probabilistic learning task with relevant information embedded in noise, combined with magnetoencephalography (MEG) recordings, we show that the learning of non-adjacent dependencies is supported by two neurally and temporally dissociable mechanisms: predictive activity and representational change. Predictive activity emerged up to 500 ms before stimulus appearance, while representational changes occurred 300-500 ms after stimulus onset and were correlated with the behavioral performance. Over the course of learning during the task, predictive activity appeared first, followed by an increase in behavioral performance and in neural representational changes. Both mechanisms primarily engaged the sensorimotor and dorsal attention networks. These findings suggest that both predictive activity and neural representational changes support the statistical learning of non-adjacent dependencies, and reveal the dynamics of these mechanisms as learning unfolds.

## Results

We recorded MEG data from 15 human participants as they performed the cued Alternative Serial Reaction Time (cASRT) task, responding to a sequence of arrows by pressing the corresponding buttons as fast and accurately as possible. Task blocks included repeating (yellow) and random (red) arrows (Fig. 1a), allowing the emergence of predictable (i.e., pattern) and unpredictable (i.e., random) elements (Fig. 1b). Participants first completed three practice blocks in which no sequence was embedded in the yellow arrows, followed by twenty learning blocks. They were informed that the yellow stimuli followed a fixed sequence in learning blocks and were instructed to identify it and optimize their performance using this knowledge.

**Fig. 1.**
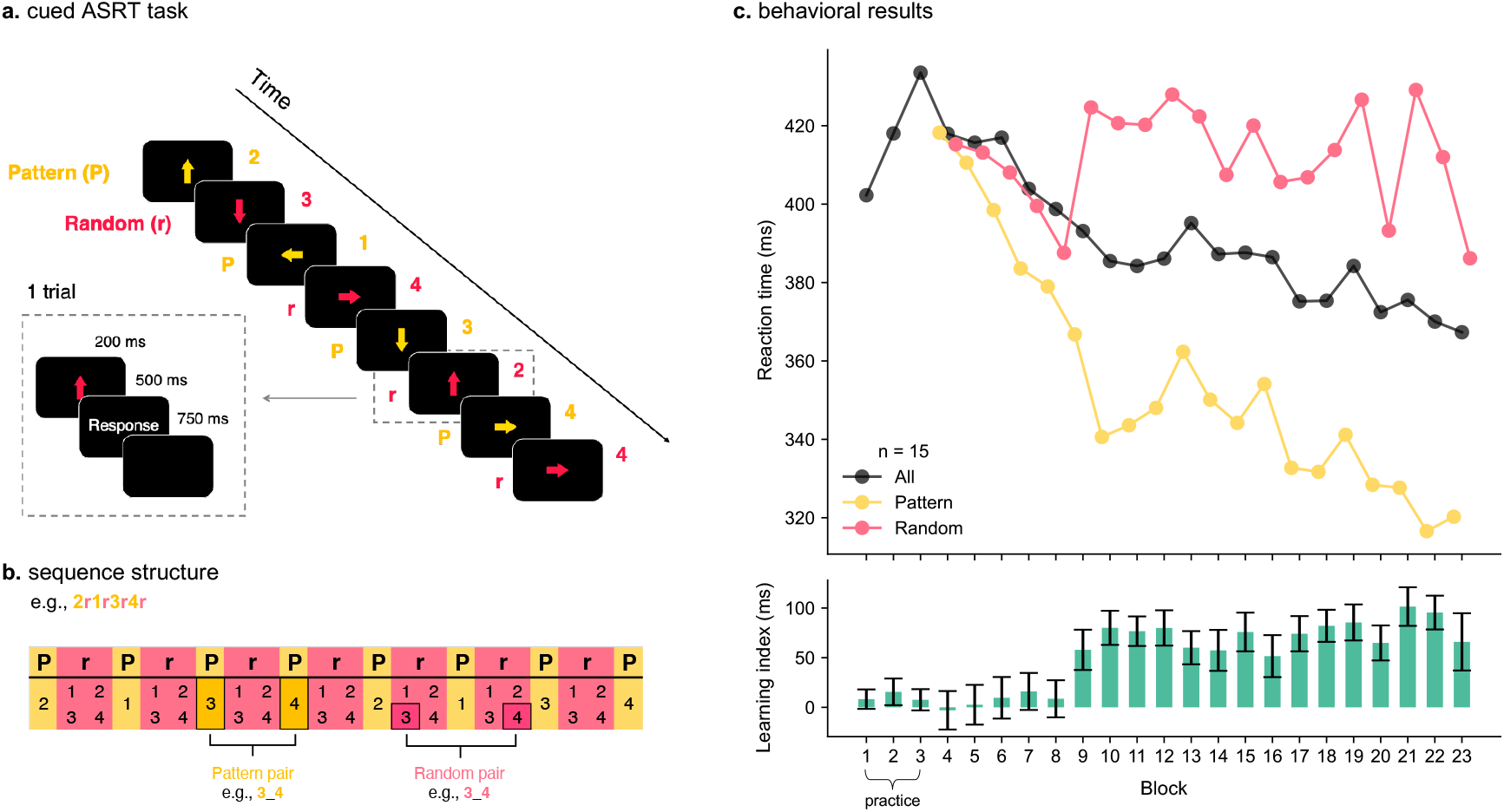
The cued Alternating Serial Reaction Time (cASRT) task leads to significant statistical learning of non-adjacent dependencies. **a** In the cASRT task, an arrow stimulus appears at the center of the screen, alternating between predictable pattern trials (yellow) and unpredictable random trials (red). Each arrow is displayed for 200 ms, followed by a 500 ms response window. The inter-trial interval is 750 ms. **b** The alternating sequence structure with numbers denoting spatial directions (1 = left, 2 = up, 3 = down, 4 = right). Statistical learning is measured as the reaction time (RT) difference in ms between pattern and random pairs. **c** Participants’ (*n* = 15) RTs for pattern and random stimuli. Colored lines show the average RTs across participants per block and trial type. Each block comprises 85 trials each of cASRT. Practice block pattern trials do not follow any sequence. The bottom panel shows the learning index (ms) per block, defined as the RT difference between random and pattern pairs. Error bars indicate the SEM. Participants were significantly faster for pattern than random stimuli, and this difference increased across blocks, confirming statistical learning. Source data are provided as a Source Data file.

Reaction time (RT) averaged over all trials for each block decreased over time (Fig. 1c) (main effect of block: F_19, 228_ = 8.46, *p* < .0001 Greenhouse-Geisser corrected), demonstrating that participants became faster independently of trial conditions. Average RT was smaller for pattern pairs than random pairs (main effect of trial condition: F_1,12_ = 22.62, *p* < .001 Greenhouse-Geisser corrected), demonstrating that participants became faster for pattern pairs compared to random pairs. Finally, the difference in RT between random and pattern pairs (i.e., learning index) increased over blocks (interaction of block and trial condition: F_19, 228_ = 2.88, *p* = .02 Greenhouse-Geisser corrected). These results confirm that statistical learning has occurred during the task. A learning index was calculated for each block as the difference in average RT between random and pattern pairs to better visualize the statistical learning across blocks.

### Information about the upcoming stimulus is present before its appearance

Predictive processing is a candidate mechanism that may underpin statistical learning of non-adjacent dependencies, either through top-down signals from higher cognitive regions or by pre-activating neural ensembles engaged during stimulus perception, even before its appearance. To investigate predictive activity, we used a Logistic Regression classifier with Leave-one-Block-out (LoBo) cross-validation strategy on both random and pattern trials and the temporal generalization^29^ method to capture neural representations and investigate if they generalize before the stimulus appearance. This analysis showed neural representations of the stimulus before the stimulus onset for pattern trials (*p* < .01, cluster-based permutation *t*-test; Fig. 2a) but not for random trials (all *p* > .01, cluster-based permutation *t*-test; Fig. 2b). When contrasted (i.e., Time generalization matrix of pattern trials – matrix of random trials), we found a significant cluster reflecting the neural representation of the upcoming stimulus, emerging ∼500 ms before its onset and building up to the stimulus appearance (Fig. 2b, yellow and black rectangles). A within-participant correlation between predictive activity and the learning index across blocks, followed by a cluster-based permutation test on the resulting Spearman correlations at the group level, revealed no significant cluster (all *p* > .05, cluster-based permutation *t*-test; Fig 2d). In summary, we found a significant predictive activity of the upcoming stimulus around 500 ms before its appearance during highly predictive trials.

**Fig. 2.**
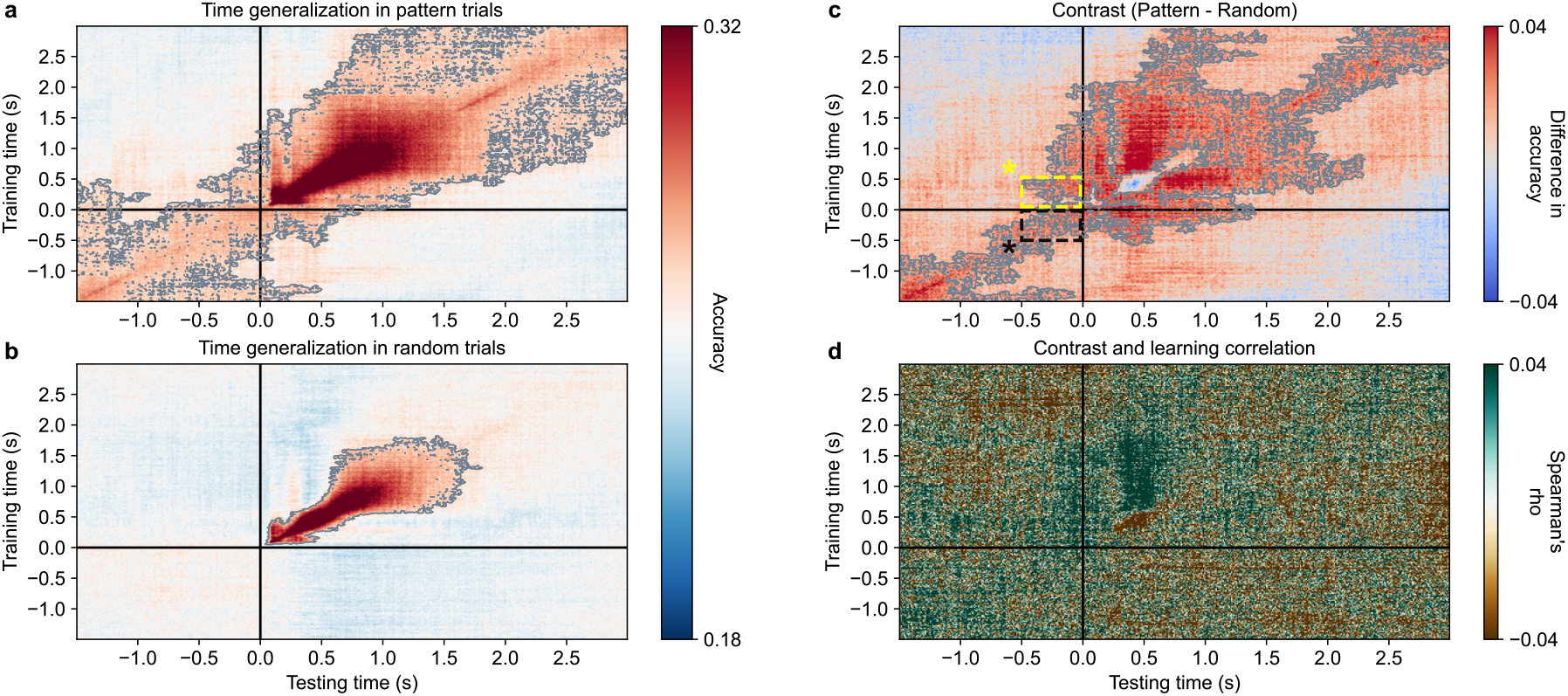
Temporal generalization of pattern classification: a predictive activity is observed before expected stimuli in pattern trials. **a** Time generalization of spatial arrows during pattern trials. Black lines indicate the stimulus onset (0 s). Statistically significant temporal clusters (cluster-based permutation *t*-test, two-sided, *p* < .01, cluster-corrected) are contoured in grey. Neural representations were found both pre- and post-stimulus. **b** Same as **a** but for random trials. Neural representations were found post-stimulus only. **c** The contrast between pattern and random trials time generalization matrices returned significant clusters in the inter-trial interval before the stimulus onset demonstrating predictive activity in pattern trials. The black dashed rectangle area represents general predictive activity, while the yellow dashed rectangle area corresponds to predictive activity that resembles the perceptual or motor activity evoked by the same stimulus. **d** The Spearman correlation between the contrast matrix and learning index returned no significant clusters (cluster-based permutation *t*-test, two-sided, all *p* > .05, cluster-corrected). Data are mean across *n* = 15 participants. \**p* < .05. Source data are provided as a Source Data file.

### Distributed predictive activity in the sensorimotor, dorsal attention, central executive, and default mode networks

To identify the brain spatial localization of the predictive activity effect, we performed source reconstruction of our MEG data using beamforming^30,31^ and investigated ten regions of interest (ROIs) comprising seven networks^32^ and three subcortical regions: visual, sensorimotor, dorsal attention, salience, limbic, central executive, default mode networks, the hippocampus, thalamus, and the cerebellum. Sensor-space analyses revealed that the effect was strongest and statistically significant mainly along the diagonal preceding stimulus onset (Fig. 2c, black rectangle), reflecting training and testing at identical pre-stimulus time points^33^. Only these segments were considered for the source-space analyses (see Fig. S1 for full temporal generalization matrices). In both pattern and random trials, decoding in source-space revealed above-chance accuracy during post-stimulus onset periods in all ROIs (Fig. 3a & 3b). In pattern trials, above-chance decoding performance during pre-stimulus onset periods was found in the sensorimotor, dorsal attention, central executive, default mode, the hippocampus, and cerebellar networks (all *p* < .0001, GAMM; Fig. 3a). We then computed the contrast (i.e., the difference between pattern and random trials decoding performance), which displayed significant clusters in the pre-stimulus period in the sensorimotor (*p* < .0001), dorsal attention (*p* < .0001, GAMM), central executive (*p* < .01, GAMM), and default mode (*p* < .05, GAMM) networks (Fig. 3c) with the sensorimotor and dorsal attention networks exhibiting the largest clusters. To observe whether predictive activity in specific regions was correlated with behavioral performance, we performed a within-participant correlation analysis across blocks followed by generalized additive modelling. No significant correlation with the learning index was found in any of the ROIs (see Fig. S2). Full temporal generalization was conducted in source space and revealed no significant clusters (see Fig. S1) corresponding to predictive activity resembling the perceptual or motor activity evoked by the same stimulus (i.e., when classifiers are trained after the stimulus onset, yellow dashed rectangle in the Fig. 2c).

**Fig. 3.**
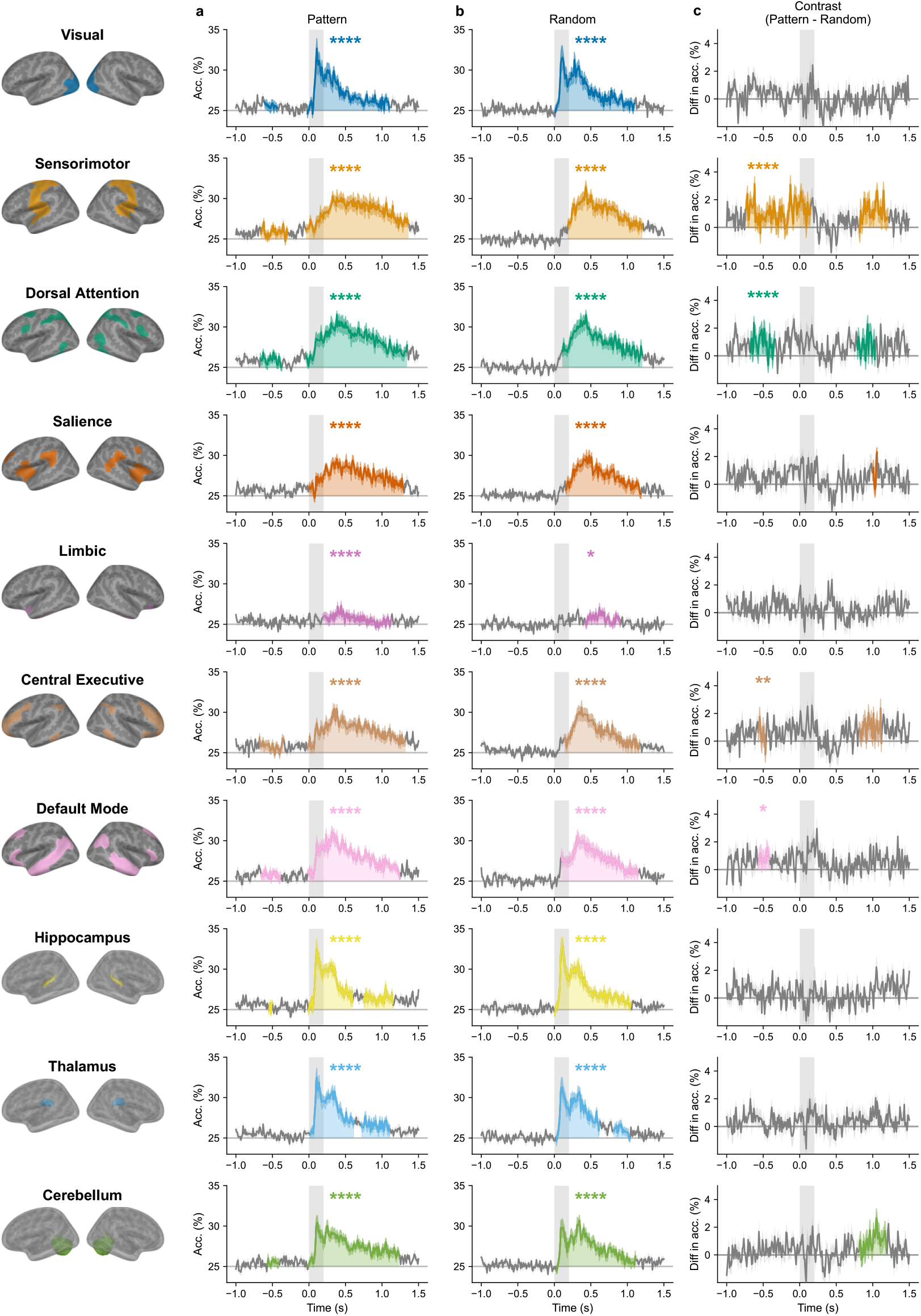
Source space decoding: a distributed predictive activity across sensorimotor, dorsal attention, central executive, and default mode networks. **a** Decoding performance of spatial arrows in pattern trials across all ROIs, using a logistic regression classifier with leave-one-block-out cross-validation, averaged across participants. The grey rectangle indicates stimulus duration (200 ms). Above-chance decoding was found post-stimulus in all ROIs, and both pre- and post-stimulus in the visual, sensorimotor, dorsal attention, central executive, default mode networks, the hippocampus, and the cerebellum. **b** Same as **a** but with random trials. Above-chance decoding was found post-stimulus only across all ROIs. **c** The contrast between pattern and random trials decoding performance. Significant generalized additive mixed model (GAMM) clusters were observed in the sensorimotor (*p* < .0001, Cohen’s *d* = 1.30), dorsal attention (*p* < .0001, Cohen’s *d* = 1.19), central executive (*p* = .0040, Cohen’s *d* = .44) and default mode (*p* = .0436, Cohen’s *d* = .62) networks. Across all panels, color-filled areas indicate timepoints at which GAMM-estimated mean differs significantly from 0 (two-sided, *p* < .05). Asterisks indicate overall GAMM significance against a null hypothesis of a constant zero temporal trend (\*\*\*\**p* < .0001, \*\**p* < .01, \**p* < .05; Holm-corrected across networks). Data are mean ± SEM across n = 15 participants. Source data are provided as a Source Data file.

In summary, our findings show a distributed predictive activity building up to the stimulus onset across the sensorimotor, dorsal attention, central executive, and default mode networks.

### Late representational change after stimulus onset at the brain level correlates with behavioral performance

Representational change, the increase in similarity of neural activity patterns between associated stimuli, is another candidate mechanism that may support statistical learning of non-adjacent dependencies. To investigate this, we conducted representational similarity analysis (RSA) using a LoBo cross-validated Mahalanobis distance (cvMD)^34,35^ between elements of both pattern and random pairs (Fig. 4a) to determine whether pairs appearing within the sequence (i.e., pattern yellow trials) exhibited greater neural similarity than the same pairs in random (red) trials. The distance within pattern pairs was subtracted from that of the random (see formula in *Methods* section). This yielded a similarity index that reflects the similarity within pattern pairs relative to the similarity within random pairs (Fig. 4b). The time course of this similarity index revealed a significant positive cluster ranging from 0.34 to 0.48 s post-stimulus onset (*p* < .001, GAMM). To investigate the link between learning and such representational change, we calculated, within participants, the correlation coefficient between the similarity index and the learning index across learning blocks (Fig. 4c). The Spearman correlation coefficient across participants revealed significant clusters from 0.30 to 0.53 s post-stimulus onset (*p* < .05, GAMM). To better visualize this relationship, we computed the average similarity index within the 0.34-0.48 s time window for each participant in each block and plotted it against the learning index for each participant (Fig. 4d). The correlation included practice blocks for both the similarity and learning indices; when these blocks were excluded, the positive correlations remained significant (see Fig. S3).

**Fig. 4.**
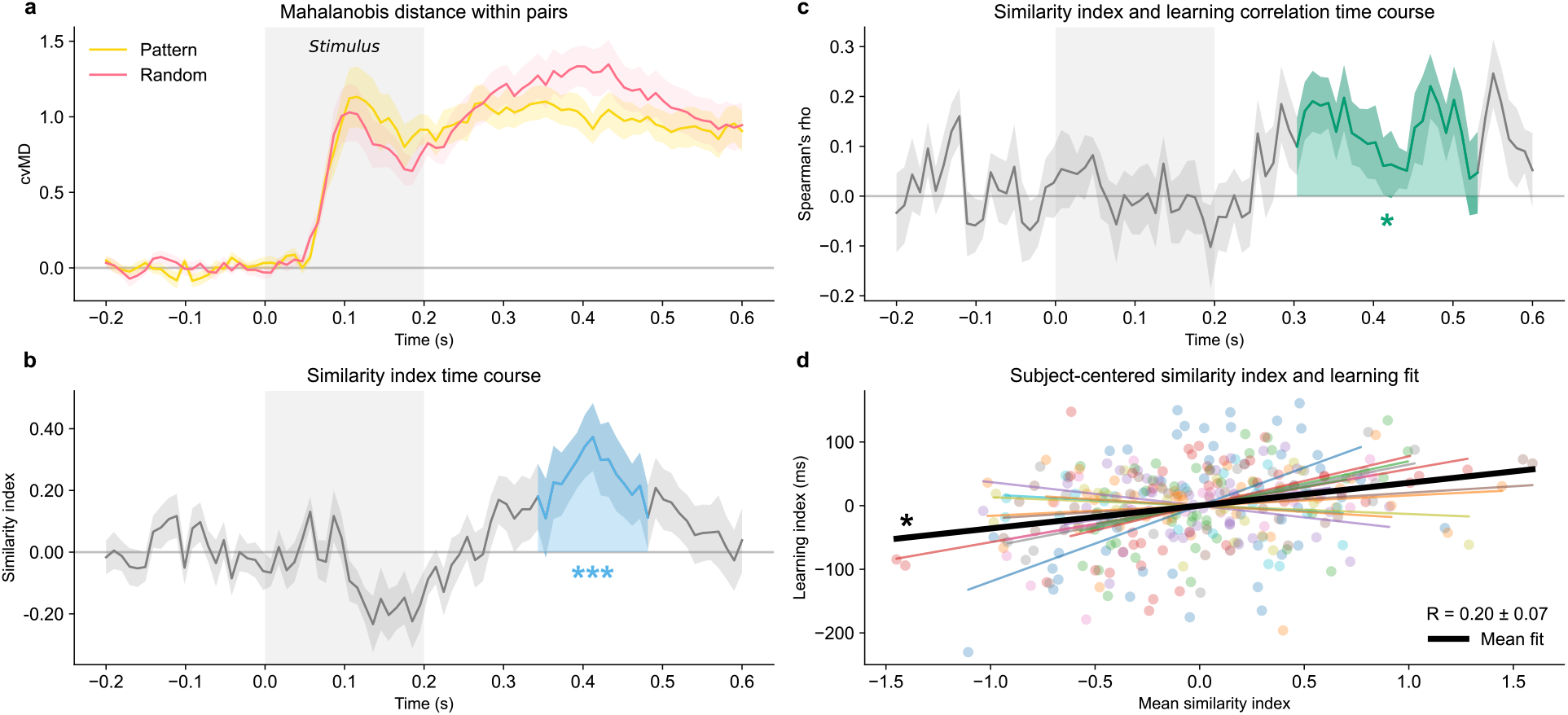
Representational similarity analysis (RSA): late representational change positively correlates with learning. **a** Time course of the leave-one-block-out cross-validated Mahalanobis distance (cvMD) for pattern and random pairs, averaged across participants. The grey rectangle indicates stimulus duration (200 ms). Shaded areas indicate the SEM. **b** Time course of the similarity index, averaged across participants. Solid lines and shaded areas indicate the mean and SEM, respectively. Elements within pattern pairs evoked more similar brain activities than elements within random pairs from 0.34 to 0.48 s post-stimulus (GAMM cluster, *p* = .0007, Cohen’s *d* = .63, two-sided). **c** Time course of Spearman’s ρ correlation between the similarity and the learning indices across blocks. Solid lines and shaded areas indicate the mean and the SEM, respectively. The similarity index positively correlates with learning from 0.30 to 0.53 s post-stimulus (GAMM cluster, *p* = .0203, two-sided). **d** Relationship between the mean representational change and the learning index across blocks. Mean representational change was computed by averaging the similarity index between 0.34 and 0.48 s for each block and each participant. Each colored line shows the linear fit between similarity and learning indices for a single participant across all blocks. The bold black line shows the mean fit across participants. Within-participant Spearman ρ values are significantly different from zero (*p* = .0197, two-sided one-sample *t*-test), confirming that increased learning is associated with greater representational change. In **b** and **c**, color-filled areas indicate timepoints at which GAMM-estimated mean differs significantly from zero (two-sided, *p* < .05), and asterisks indicate overall GAMM significance against a null hypothesis of a constant zero temporal trend (\*\*\**p* < .001, \**p* < .05). Data are mean ± SEM across *n* = 15 participants. Source data are provided as a Source Data file.

Overall, these findings demonstrate that the brain updates its representations of pattern pairs relative to random pairs by increasing the similarity of non-adjacent elements within pattern pairs compared with the same elements when presented in non-informative random positions. The positive correlation with learning suggests that representational changes are linked to the learning process, either supporting it or being shaped by it.

### Distributed representational change across the visual, dorsal attention, salience, central executive networks, and the cerebellum

To investigate representational change at the source level, we employed the same source reconstruction pipeline as for testing predictive activity. RSA was performed in each ROI using the same LoBo cvMD distance measure (Fig. 5a) as in the previous sensor-space analysis. The visual (*p* < .001, GAMM), dorsal attention (*p* < .0001, GAMM), salience (*p* < .05, GAMM), central executive (*p* < .05, GAMM), and cerebellar (*p* < .0001, GAMM) networks returned significant clusters around 400 ms post-stimulus onset, with the dorsal attention network exhibiting the largest cluster and the visual network the shortest (Fig. 5b). The time course correlation between the similarity index with learning revealed a brief significant cluster in the dorsal attention network around 400 ms post-stimulus onset (*p* < .05; Fig. 5c). When practice blocks were excluded, no significant correlations remained in any of the ROIs, except in the dorsal attention network before stimulus onset, which is likely a false positive as it only reached the lowest significance threshold (*p* < .05, GAMM) and its temporal position (beginning before stimulus onset) is inconsistent with a possible change in representational similarity (see Fig. S4).

**Fig. 5.**
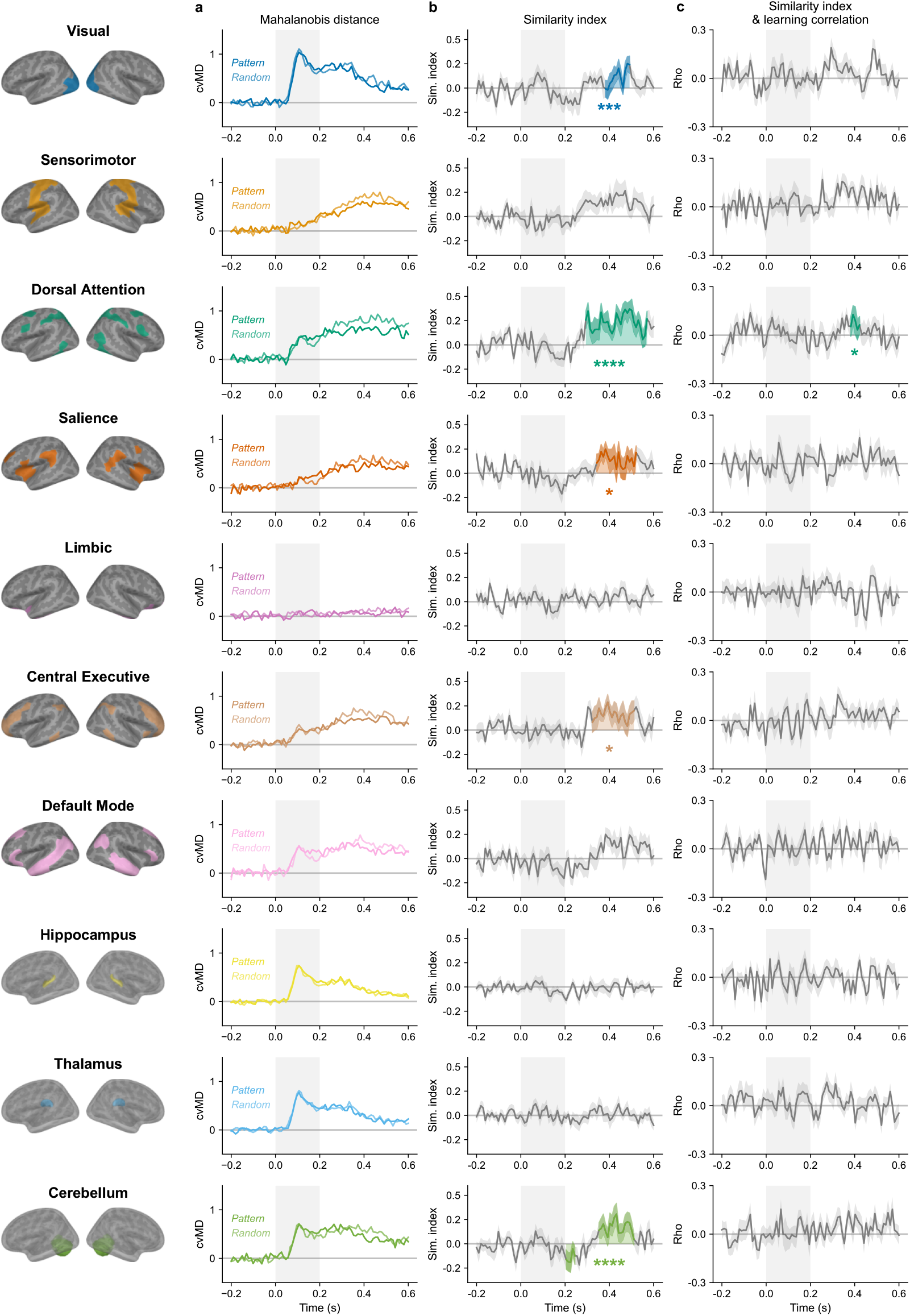
Source space RSA: distributed representational change in the visual, dorsal attention, salience, central executive networks, and the cerebellum. **a** Leave-one-block-out cross-validated Mahalanobis distance for pattern and random elements averaged across participants. The grey rectangle indicates stimulus duration (200 ms). Shaded areas indicate the SEM. **b** Similarity index time course across ROIs, averaged across participants. Significant GAMM clusters of representational change were found in the visual (*p* = .0007, Cohen’s *d* = .34), dorsal attention (*p* < .0001, Cohen’s *d* = 1.02), salience (*p* = .0161, Cohen’s *d* = .22), central executive (*p* = .0313, Cohen’s *d* = .53), and cerebellar (*p* < .0001, Cohen’s *d* = .73) networks. Shaded areas indicate the SEM. **c** Spearman’s ρ correlation time course between the similarity and learning indices across learning blocks. Only the dorsal attention network showed a significant GAMM cluster (*p* = .0165). In **b** and **c**, color-filled areas indicate timepoints at which GAMM-estimated mean differs significantly from 0 (two-sided, *p* < .05). Asterisks indicate overall GAMM significance against a null hypothesis of a constant zero temporal trend (\*\*\*\**p* < .0001, \*\*\**p* < .001, \**p* < .05; Holm-corrected across networks). Data are mean ± SEM across *n* = 15 participants. Source data are provided as a Source Data file.

In summary, these findings indicate that the initial representational change observed at the brain level is distributed across the visual, dorsal attention, salience, central executive, and cerebellar networks at the source level. No strong association was found between the magnitude of learning and representational change across these networks, except a small significant cluster for the dorsal attention network.

### Non-linear temporal modelling on behavior, predictive activity, and representational change reveals distinct temporal dynamics across blocks and ROIs

In order to track changes in representations and predictions over the course of learning, we applied non-linear, generalized additive mixed models (GAMMs) to the block-wise time courses of behavioral performance, predictive activity, and representational changes (Fig. 6). First, we fitted one distinct model to each measure. We found that behavioral performance showed a statistically significant increase from block 6 to block 10 before reaching a plateau, and was significantly above 0 from block 8 onward (*p* < .0001, GAMM). Regarding neural measures, predictive activity appeared to rise from the very first block of practice until block 6 before reaching a plateau and was significantly above 0 from block 4 onward (*p* < .0001, GAMM). Representational change increased throughout the behavioral task (block 1 to 23) and was significantly above 0 from block 9 until the end of experiment (*p* < .05, GAMM). While the statistical findings from the individual models suggest that behavioral performance, predictive activity, and representational change have different temporal dynamics as learning unfolded, we collected more direct evidence from models of pairwise differences between the 3 measures. These models were fitted on standardized data so that all measures were brought to the same arbitrary unit scale. We found a significant main effect of *block* for the difference between behavioral performance and predictive activity (*p* = .02, GAMM), which was driven by a significant difference between blocks 4 and 9. This result confirms that predictive activity manifests several blocks before the improvement of behavioral performance. On the other hand, the temporal dynamics of representational similarity was not statistically distinguishable from that of the other measures.

**Fig. 6.**
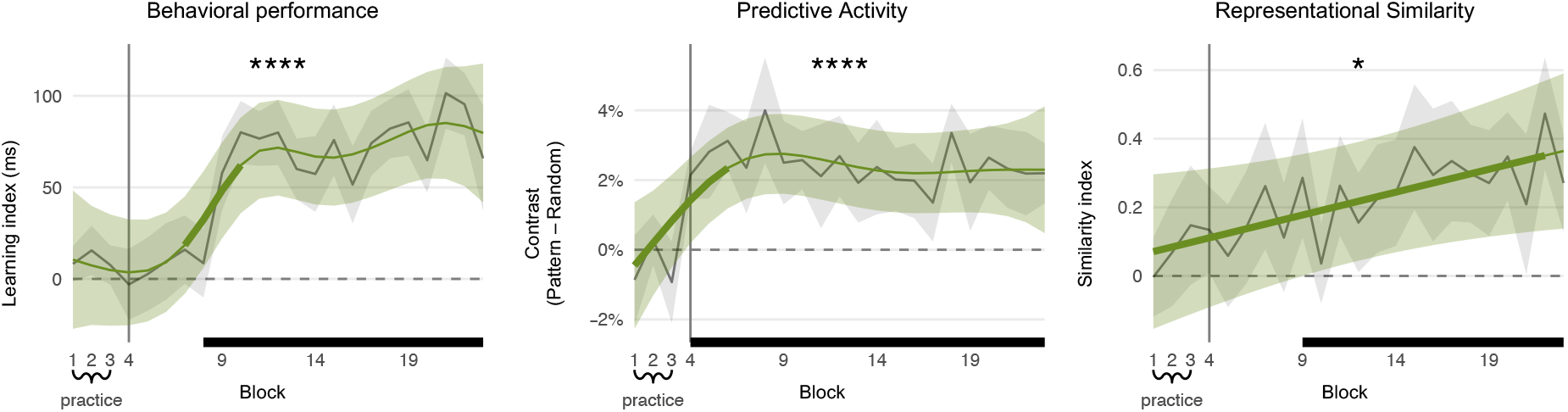
Behavior, predictive activity, and representational change have different temporal dynamics across blocks. Vertical grey lines separate practice from learning blocks. Grey lines and ribbons show the group mean ± SEM. Green lines and ribbons show the GAMM-estimated marginal means and 95% CI. Horizontal black segments mark blocks whose GAMM-estimated mean differs significantly from 0 (two-sided, *p* < .05). Thicker portions of the green smooth mark blocks with a significant temporal slope (GAMM derivative test, two-sided, *p* < .05). Asterisks indicate overall GAMM significance against a null hypothesis of a constant zero temporal trend (\*\*\*\**p* < .0001, \**p* < .05; Holm-corrected for metrics). Behavioral performance is significantly above 0 from block 8 (*p* < .0001, GAMM), predictive activity from block 4 (*p* < .0001, GAMM), and representational change from block 9 (*p* = .0468, GAMM). Data are mean ± SEM across *n* = 15 participants. Source data are provided as a Source Data file.

We conducted the same analysis with mean effects of representational change and predictive activity in source space across all our ROIs (Fig. 7). Predictive activity was significantly above 0 in the sensorimotor and dorsal attention networks from block 1 onward, and salience network from block 1 to block 19 (all *p* < .001, GAMM). Representational change was significantly above 0 in the sensorimotor network from block 12 onward (*p* < .01, GAMM) and the dorsal attention network from block 8 onward (*p* < .0001, GAMM). In summary, our results suggest that predictive activity and representational change have different temporal dynamics mainly distributed across the sensorimotor and dorsal attention networks.

**Fig. 7.**
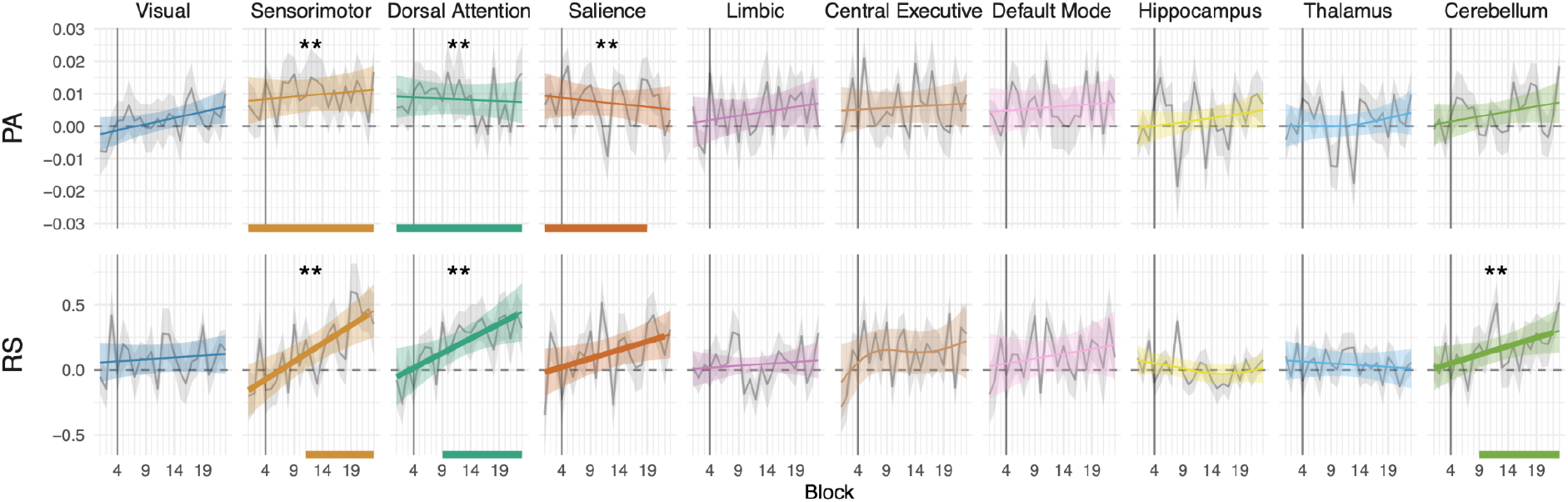
Distributed predictive activity and representational change effects across blocks in sensorimotor and attentional networks. Top row: predictive activity (PA). Bottom row: representational similarity (RS). Vertical grey lines separate practice from learning blocks. Grey lines and ribbons show the group mean ± SEM. Colored lines and ribbons show the GAMM-estimated marginal means and 95% CI. Horizontal colored segments mark individual blocks whose GAMM-estimated mean differs significantly from 0 (two-sided, *p* < .05). Thicker portions of the colored smooth mark individual blocks with a significant temporal slope (GAMM derivative test, two-sided, *p* < .05). Asterisks indicate overall GAMM significance against a null hypothesis of a constant zero temporal trend (\*\**p* < .01; Holm-corrected across networks). Predictive activity is significantly above 0 from block 1 in the sensorimotor (*p* = .0021, GAMM), dorsal attention (*p* = .0041, GAMM), and salience (*p* = .0057, GAMM) networks. Representational change is significantly above 0 in the sensorimotor network from block 11 (*p* = .0027, GAMM), in the dorsal attention from block 9 (*p* = .0027, GAMM), and in the cerebellum from block 9 (*p* = .0089, GAMM). Data are mean ± SEM across *n* = 15 participants. Source data are provided as a Source Data file.

## Discussion

To adapt to a dynamic world, the human brain must extract stable regularities from noisy environments, a process known as statistical learning^1,3^. While the neural basis of learning simple adjacent dependencies has been explored^14–16^, how the brain learns complex, non-adjacent patterns separated by irrelevant information remains poorly understood. Here, using MEG during a visuomotor task, we demonstrated that learning non-adjacent dependencies is supported by two neurally and temporally dissociable mechanisms: predictive activity and representational change. We first reveal an early emergence of predictive activity (Fig. 6), where neural patterns specific to expected stimuli appear up to 500 ms prior to their onset (Fig. 2) across a distributed network including sensorimotor, dorsal attention, central executive, and default mode networks (Fig. 3). Second, we found a more gradual representational change (Fig. 6), where the neural patterns evoked by linked, non-adjacent stimuli become more similar. This representational shift occurs between ∼300 to 500 ms after stimulus onset (Fig. 4), and is distributed across large-scale networks that include the visual, sensorimotor, dorsal attention, salience, central executive networks, and the cerebellum (Fig. 5 & Fig. 7). Notably, our temporal modelling across task blocks reveals distinct dynamics between the two neural mechanisms: predictive activity emerged first, followed by behavioral improvements, and neural representational changes that continued to increase throughout the task (Fig. 6). These results suggest that fast predictions precede initial performance gains while slower representational plasticity solidifies learned structures.

Extensive evidence supports that the brain continuously attempts to match incoming perceptual input through predictive activity shaped by previous experience. Despite neural mechanisms underlying such predictive activity being unclear, one widely accepted hypothesis is that predictive signals are transmitted through top-down pathways from higher-order to sensory regions^18,25^. We used the temporally generalized multivariate pattern classification method^29^ to identify neural predictive activity. In sensor space, when classifiers were both trained and tested within the pre-stimulus period, we observed above-chance decoding performance before the onset of the predictable stimulus only (Fig. 2a-b). This observation suggests that the brain pre-activates a stimulus-specific activity before the onset of the predictable (pattern) stimulus. When classifiers are trained and tested before the onset of the stimulus (Fig. 2c, black dashed rectangle), such predictive activity is not necessarily similar to the activity during the perception of the same stimulus and could be attributed to predictions in high-level regions (e.g., frontal lobe). At the source-level, the sensorimotor, dorsal attention, central executive, and default mode networks revealed such predictive activity within the 500 ms pre-stimulus period (Fig. 3).

By contrast, above-chance decoding performance when classifiers are trained after stimulus onset but tested before (i.e., the yellow dashed rectangle in Fig. 2c) would represent predictive activity resembling the perceptual or motor activity evoked by the same stimulus^18^. However, we found no evidence for such a pre-activation effect at the source level (see Fig. S1). In our sensor-level results, both interpretations remain possible but should be considered with caution due to potential temporal leakage within the larger significant cluster^36^. Similarly, in sensor space, the significant cluster in pattern trials corresponding to classifiers trained between 0 and 0.5 s and tested between −1.5 and −1 s suggests that, during the perception of a stimulus, a neural representation resembling that of the next predictable element is transiently present. This is consistent with a forward-looking predictive account, whereby the current stimulus triggers a rapid pre-activation of the upcoming one^14^. However, as this effect did not reach significance in the contrast between pattern and random trials nor in source space analyses, its interpretation remains speculative and warrants investigation in future studies. Pre-activation of neural ensembles prior to stimulus onset, showing activity similar to that during the actual perception of the same stimulus, has already been proposed as a candidate mechanism for neural prediction^24,25^. We have challenged previous work in humans investigating this mechanism on methodological grounds, as the high physical similarity between two consecutive stimuli in previous experimental designs can produce spurious effects that confound genuine prediction^26^. The structure of our cASRT task, where each preceding (*n-1*) trial is always uninformative of the current (*n*) trial, allows us to isolate anticipatory neural activity from such confounds, despite physical similarity of the linked stimuli. Yet, because our paradigm is fully deterministic between two pattern trials without any jitter, it is theoretically possible to confound memory traces from preceding trials for predictive activity. Several lines of evidence suggest otherwise. Indeed, predictive activity is not present during the practice blocks (Fig. 6) and source-space decoding is not continuously significant before stimulus onset (Fig. 3). Additionally, we conducted an empirical control analysis^26^ by constructing continuous raw datasets by re-ordering random trials to match the sequence structure (see Fig. S5). This analysis failed at replicating significant decoding performances during the pre-stimulus period (Fig. 2), further confirming that the observed neural patterns represent active predictive activity rather than a residual memory trace. Moreover, it is important to note that while the predictable elements in our cASRT task would technically form an adjacent deterministic sequence if the random trials were removed, they remain strictly non-adjacent from a cognitive perspective. The sensorimotor and attentional networks must disengage from the underlying sequence to process the unpredictable intervening event. Therefore, extracting the dependency requires bridging a cognitive gap, closely mimicking the demands of non-adjacent dependency processing found in natural environments, such as language (e.g., linking a subject and a verb across a relative clause).

Predictive coding theories do not necessarily posit that predictive activity reflects a literal pre-activation of sensory templates^37–39^ (i.e., activity resembling the perception of the same stimulus). Instead, higher-order regions may prepare relevant neuronal populations through gain control mechanisms^39^. In this framework, the pre-stimulus signal reflects a shift in synaptic sensitivity of specific populations, rather than a generative re-activation of the upcoming stimulus representation. Our results are more consistent with this view, and relate to findings showing that memory representations do not share the same neural activity patterns as those evoked during perception of the same content^40^. More broadly, predictive coding theories^21,41,42^ posit that the brain seeks to infer the causes of sensory inputs by minimizing surprise, optimizing prediction and interaction with the environment. Press et al. (2020) proposed the opposing process theory (OPT) to reconcile conflicting findings showing that expected stimuli can be either perceptually enhanced or suppressed depending on the context^37^. They suggest that both processes operate sequentially, with the perception initially biased towards expected stimuli (i.e., sharpening), before switching to prioritize unexpected stimuli (i.e., dampening). Consistent with this view, we observed higher decoding performance for pattern than random stimuli at early post-stimulus latencies (∼0.15 to 0.30 s after stimulus onset), followed by higher decoding for random than pattern stimuli at later latencies (∼0.40 s) (see Fig. S6a-b), consistent with recent empirical evidence^38^. Furthermore, these effects were also observable across blocks with the sharpening effect showing a rapid rise peaking at block 10 then plateauing, and the dampening raising more slowly from block 10 onward (see Fig. S6c). Non-linear temporal modelling revealed a significant cluster during the sharpening window only at the single trial level (see Fig S6b). The dynamic across the experiment (block-level, Fig. S6d) suggests that learned expectations first sharpen and then dampen neural representations of predictable stimuli but warrants further investigation with larger cohorts.

Schapiro et al.^14,16^ evidenced representational change as a mechanism of statistical learning of adjacent dependencies using a passive exposure task. Their fMRI study revealed the hippocampus and the medial temporal lobe as areas that support this mechanism. Our findings extend this work by showing that representational change also supports statistical learning of non-adjacent dependencies, indicating that this mechanism remains present when regularities are embedded within noisy information. Moreover, representational change positively correlates with behavioral performance (Fig. 2d). Our findings differ from Schapiro et al.^14^ in two ways. First, we did not find evidence of representational change in the hippocampus (Fig. 5). This difference may arise from the difficulty of resolving subcortical signals in MEG source space, where signal-to-noise ratios are generally low, or from the use of a different statistical learning task involving motor responses. Second, in our explicit task, participants were aware of the presence of a sequence within the stimulus stream, which facilitated learning^7,13^ and may have accelerated the emergence of representational changes. Schapiro et al. reported that representational change emerged only after an hour of stimuli exposure^16^ while, in our study, the effect is present after about 18 min (from block 9) of sequence exposure (Fig. 6). The quicker emergence of the effect may be due to the inclusion of a motor response, which likely increased participants’ engagement and active involvement in the task compared to the passive exposure task used in Schapiro et al.^14,16^.

We also build upon previous temporally resolved RSA studies^12,15^, that identified representational changes for predictable auditory stimuli, extending these findings to visual perception and the learning of non-adjacent dependencies. Our results contrast with a late effect (i.e., 300 to 500 ms post-stimulus onset) compared to an early effect (i.e., around 120 ms post-stimulus onset) of previous studies, which could also be attributed to our experimental paradigm that incorporates a motor response. Further to this, our results of representational change show a distributed mechanism across various large-scale networks rather than a localized one. This is in line with previous works^12^ suggesting that statistical learning engages distributed cortical networks involved in the parallel encoding of both low (e.g., transitional probabilities) and high-level statistical features (e.g., sequence order and identity).

Quantifying effects of behavioral performance, predictive activity, and representational change across blocks revealed distinct dynamics for each process (Fig. 6). Predictive activity emerged first, becoming statistically significant right from the first block of learning and quickly reaching a plateau. Such rapid emergence of predictive activity from the first learning block may reflect the explicit nature of the task, in which participants are informed of the presence of a hidden sequence prior to learning. However, improvement in behavioral performance was detectable only several blocks later. Similarly, the explicit knowledge of the sequence was high (>60%) from the first block of learning but reached a plateau more slowly (see Fig. S7). This temporal dissociation, added to the absence of evidence of a correlation between predictive activity and behavior (Fig. 4d) suggests that predictive activity and explicit knowledge do not necessarily or immediately translate to behavioral change. On the other hand, representational similarity appeared to increase slowly but continuously across the entire experiment. However, statistical models could not establish which of the two candidate mechanisms (representational change or neural predictions) preceded the other, if any. One hypothesis compatible with the observed pattern of temporal dynamics is that predictive activity drives representational change, which in turn supports behavioral improvement. Slower representational change, particularly in attentional networks, may provide long-term memory consolidation, explaining why representational similarity continues to rise even after behavioral performance plateaus. Together, these findings support the engagement of large-scale neural interactions in predictive processing and representational change in response to environmental statistical regularities^15^. Historically, consolidation research focused on extended rest periods, such as hours-long delays or sleep. However, memory stabilization is now known to occur over minutes or even seconds^41^. Procedural and sequential skills often consolidate during the brief breaks between practice blocks, a process termed micro-offline or ultra-fast consolidation^41–44^. The continuous build-up of representational similarity we observed across blocks could reflect such rapid consolidation. During inter-block rests, the brain may rehearse task-relevant associations, possibly through neural replay^45^, gradually restructuring neural geometry within the dorsal attention and sensorimotor networks and transforming transient experiences into durable representations. Future studies will be needed to untangle the precise mechanistic relationship between micro-scale online and offline behavioral gains, predictive neural activity, representational change, and their causal relationships.

The role of attentional networks has been strongly linked to non-adjacent dependency processing involving top-down signals from cognitive control networks to inhibit attention toward intervening items and redirect it toward the relevant dependencies^10^. Our results in representational change and predictive activity corroborate these previous studies. Indeed, dorsal attention is involved in both predictive activity (Fig. 5 & 7) and representational change (Fig. 3 & 7). The sensorimotor network exhibits predictive activity (Fig. 5 & Fig. 7), coherent with motor preparation activity before a predictable stimulus. Although less evident in representational similarity analyses (Fig. 3), the sensorimotor network shows a significant build-up when tracked across blocks (Fig. 7). The cerebellum contributed reliably to representational change but not to predictive activity, suggesting that, in the context of our task, its role lies in updating and consolidating associative mappings rather than pre-activating representations, as is often proposed^46^. This aligns with models of the cerebellum as an error-driven learning system that refines internal models across repetitions^47^ (i.e., representational change in the cerebellum occurs within the motor response time range), thereby restructuring and stabilizing representational geometry without directly generating anticipatory activation^48^. It also converges with evidence for cerebellar involvement in working memory^48,49^ and, more broadly, in language processing^48,50^ which is strongly linked to statistical learning^3,6^. The visual network is also involved in representational change (Fig. 5). Interestingly, its contribution occurs in the range of the response time window (i.e., 300-500 ms). This particular time range suggests that statistical learning promotes post-perceptual transient representational change. The late visual representational change indicates first that the visual representations are reactivated shortly before or during the motor response, and, second that, these visual patterns become more similar in the context of response selection. Findling et al.^51^ found that the neural representations of prior expectations about the environment in mice during a decision-making task engaged both cortical and subcortical areas, spanning from early sensory to higher-order areas. Our results echo these findings in the context of a visuomotor task with humans. While we primarily observed predictive activity in the sensorimotor and dorsal attentional networks, such predictive activity was also present in the central executive and default mode networks. This suggests that the central executive and the default mode networks may play a supportive role in predictive activity^18–20,52^ by generating top-down priors to attentional and sensorimotor networks in anticipation of upcoming stimuli. In summary, we observed that both representational change and predictive activity engage distributed brain networks. Two categories of contributors emerged: the core networks, including the dorsal attention and the sensorimotor systems, and the more transient networks, encompassing the visual, salience, central executive, default mode, and cerebellar systems. Complementing our findings, previous electrophysiological evidence shows that perceptual and motor codes contribute jointly to the predictive processes underlying sequence learning, supporting the view that probabilistic learning relies on the coordinated operation of both modality-specific and modality-independent neural representations^53^. Together, these results align with our MEG data, indicating that predictive activity and representational change arise not in isolation but through the dynamic interplay of distributed brain systems that flexibly balance top-down control and multi-level coding to optimize learning. In order to identify the contributing anatomical regions, we mapped the functional parcels of the sensorimotor and dorsal attention networks to the Destrieux anatomical atlas^54^ (see Table S1). Predictive activity is localized in the paracentral sulcus and the precuneus gyrus in sensorimotor and dorsal attention networks, respectively (see Fig. S8). Representational change is widespread across both networks, with the central sulcus for sensorimotor and the precentral superior sulcus for the dorsal attention emerging as major contributors (see Fig. S9). While these mappings provide valuable anatomical context for predictive activity and representational change, the overlap between both atlases is not exhaustive. Thus, these regions should be interpreted with caution and as most-probable anatomical drivers of the observed effects in the large-scale networks.

A question concerns the mechanisms driving representational change. Schapiro et al.^14^ proposed two possibilities: an associative account, in which the neural representations of linked stimuli converge (e.g., in pair *AB*, the representation of *A* moves closer in similarity toward *B*, and vice versa), and a predictive account, in which similarity increases because the upcoming predictable stimulus is pre-activated during perception of the current one (e.g., the representation of stimulus *B* is activated during the perception of *A*)^14^. Our task, which interleaved each pattern with a random one, enables us to disentangle these two mechanisms: representational changes emerged 300 to 500 ms after pattern stimuli onset, whereas predictive activity occurred prior to pattern stimuli, following the presentation of a random stimulus. Moreover, their distinct temporal dynamics (Fig. 6 & 7) further support the existence of two separable processes, consistent with the associative account.

One limitation of our study is the explicit nature of the task, which directs participants’ attention toward the pattern trials and the hidden sequence embedded in it. In more ecological contexts, such as natural language acquisition, no instruction is given to attend to non-adjacent dependencies (e.g., the beginning and end of a sentence): the learning occurs implicitly. Studies have shown that attentional manipulations do change sequence learning performances and strategies^55,56^ but also how non-adjacent dependencies are processed^57,58^. Our results are therefore bound to our paradigm. An important next step would be to perform the same analyses with a task involving implicit non-adjacent dependencies, or with pairs defined by graded probability distributions rather than fixed patterns, where attention is not explicitly directed to the regularities. One more limitation would be that source reconstruction in MEG data is subject to inherent spatial leakage across ROIs, particularly given our use of large-scale cortical networks and subcortical regions such as the thalamus, the hippocampus, and the cerebellum (see Fig. S10-S13). Consequently, some overlap of information between adjacent networks should be expected, which may reduce the spatial specificity of reconstructed signals and should be taken into account when interpreting the results. Another consideration is the relatively small sample size (*n* = 15). However, the high trial count per participant (∼1,700 trials) significantly increases the signal-to-noise ratio and within-participant power. Future studies with larger cohorts would be valuable to confirm these findings.

In conclusion, our work reveals that statistical learning of non-adjacent dependencies is not a monolithic process but a dynamic interplay between two mechanisms operating on different timescales. We demonstrate that the brain first employs rapid predictive activity with anticipatory signals emerging early across attentional and sensorimotor networks. This initial, fast-acting mechanism is followed by a slower, more gradual representational change, which correlates with performance gain. This temporal hierarchy, supported by interactions between attentional and sensorimotor systems, provides a mechanistic framework for how the brain extracts regularities in noisy environments, suggesting a fundamental principle whereby fast predictions pave the way for lasting representational plasticity.

## Methods

All analyses were performed using the Python programming language (v3.10.9, Python Software Foundation, https://www.python.org/) with main packages including MNE-Python^59^ and Scikit-learn^60^, and R language (version 4.3.2).

### Participants

Participants were recruited in France through promotional flyers. Fifteen healthy young adults took part in the experiment (10 female, 5 male; mean age = 25.8 ± 2.3 years). Handedness was measured by the Edinburgh Handedness Inventory^61^; the mean Laterality Quotient (mLQ) of the sample varied between +33.3 and +100 (−100 means complete left-handedness, 100 means complete right-handedness; mLQ = 76.0 ± 22.8). Participants had normal or corrected-to-normal vision. None of them reported a history of any neurological and/or psychiatric condition, and none of them were taking any psychoactive medication. To ensure that participants did not differ in executive function performance that would account for difference in statistical learning performance, the short form of the Counting Span test was administered before the magnetoencephalography (MEG) experimentation, for which they performed with a mean score of 3.78 (SD = 0.95) and a coefficient of variation of 0.25 indicating moderate interindividual variability but no extreme outliers. All participants provided written informed consent before enrollment and were compensated a total of 40€ for the entire experiment. The experiment was conducted in accordance with the guidelines of the Declaration of Helsinki and was approved by the “Comité de Protection des Personnes Sud Méditerranée I”.

### Behavioral task and experimental procedures

Statistical learning was measured with a cued version of the alternating serial reaction time (cASRT) task^62,63^. In this task, an arrow stimulus appears at the center of the screen and participants are instructed to press the key on a response device that corresponds to the spatial direction of the actual arrow, as quickly and accurately as possible. Participants used their index and middle finger of both their hands to respond. The cASRT task was written, delivered to the participants, and monitored using the Presentation software (v. 18.1, Neurobehavioral Systems). Visual stimuli were back-projected on a translucent screen in front of the participants. Motor responses were collected using two LUMItouch optical response keypads with two keys each (i.e., left and up response keys on the left pad; down and right keys on the right pad). The presentation of arrow stimuli follows an alternation of pattern (P) and random (r) trials (e.g., 2–r–1–r–3–r–4–r; where numbers denote the four predefined spatial directions [1 = left, 2 = up, 3 = down, 4 = right] of the arrows and r denotes randomly chosen directions out of the four possible ones) (Fig. 1a). Arrows corresponding to pattern trials were displayed in yellow and arrows corresponding to random trials in red. Participants were told that yellow arrows always follow a predefined sequence of spatial directions while red arrows always point to a randomly chosen spatial direction out of the four possible ones. After the practice blocks, during which pattern trials did not follow any sequence, they were instructed to find the hidden predefined sequence in the following blocks, defined by the yellow arrows, to improve their performance, i.e., to use this sequence information to predict the next sequence (pattern) element in order to respond faster and more accurately. However, they were not informed about the exact length of the sequence. Instructions emphasized that participants also have to respond to red (random) arrows. Out of the 24 possible permutations of the four spatial directions, 6 unique permutations were selected and assigned to each participant in a pseudorandom manner. In such a sequence, some pairs become predictable (i.e., pattern pairs) in contrast to some that stay unpredictable (i.e., random pairs) (Fig. 1b). The timing of an experimental trial was the following. The target stimulus (arrow) was presented at the center of the screen for 200 ms, then a blank screen was displayed until the participant gave a motor response but no longer than 500 ms. After a fixed delay of 750 ms following the response (indicated by a blank screen again), the next trial started (Fig. 1a). In the case of an incorrect response, an X was presented at the center of the screen for 500 ms before the 750 ms blank screen delay. If no response was recorded within 700 ms following stimulus onset (the duration encompassing both stimulus presentation and the response window) a ! appeared at the center of the screen for 500 ms, followed by a 250 ms blank screen before the onset of the next trial. Participants first performed 3 practice blocks in which pattern trials did not follow the predefined sequence yet. Then they performed 4 runs of 5 blocks with the pattern trials following the sequence for a total of 20 learning blocks. Each block included 85 trials (for a total of 1700 trials in the entire testing experimental session) after five warm-up trials consisting only of random stimuli. After each block, participants received a 3 s feedback on their mean RT and accuracy in the given block. After the feedback, explicit knowledge about the sequence was measured. Participants were instructed to continuously type the order of yellow arrows with the corresponding response keys. This sequence report lasted until participants gave 12 consecutive responses (i.e., ideally, they reported the given sequence three times). Participants were not informed that they had to provide exactly 12 responses and that the series of responses could be three repetitions of the given sequence. The explicit test was followed by a self-paced short break while participants could rest. Accuracy of the explicit test was computed per block. The entire experimental procedure lasted an average of 56 minutes including the 6 min practice blocks and the 2 min MEG recalibration between each run. Structural MRI data were acquired prior to the MEG experiment from 12 participants using a Siemens Prisma 3T scanner equipped with a 64-channel coil. High-resolution 1 mm isotropic 3D magnetization-prepared rapid gradient echo (MPRAGE) T1-weighted images were obtained (repetition time = 2400 ms; echo time = 2.98 ms; matrix size = 192 × 256 × 256).

### Behavioral analysis

Stimulus repetitions (e.g., 222, 444) and trills (e.g., 313, 424) were excluded^64^, and only correct responses were considered for further analysis. Mean RTs for pattern and random pairs were calculated separately for each participant and session. A 2 (trial conditions: pattern vs. random pairs) x 20 (learning blocks) two-way repeated-measures analysis of variance (ANOVA) was performed to compare mean RTs throughout the experiment. Greenhouse–Geisser correction was applied to the reported *p* values. For each session, the learning index was calculated with the following formula, and tested against zero with a one-sample *t*-test:

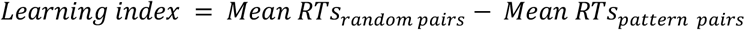

### MEG recordings and preprocessing

Neuromagnetic activity was recorded with a sampling rate of 2034.5 Hz on a whole head 248 magnetometers MEG system (4D Neuroimaging magnes 3600). The MEG apparatus was housed in a magnetically shielded room. The head position was recorded before and after each run. One electrocardiography (ECG) channel and two electrooculography (EOG) channels were also recorded and realigned during preprocessing. All trials were considered irrespective of the response given by participants. Head positions of the participants were realigned to that of the first learning run (i.e., first 5 blocks). We first applied a high pass filter above 1 Hz to the raw data. Independent component analysis (ICA) was then performed on this high-pass-filtered raw data to extract 30 components. To remove components linked to eye blinks/movements or heartbeats artifacts, we computed the Pearson correlation of each ICA component time course with each of the ECG/EOG channels. ICA components that correlated above a threshold of 0.9 with one of the ECG/EOG channels were discarded. The remaining ICA components were back-projected to the MEG sensors using the original raw data (unfiltered). This ICA-cleaned raw data was then bandpass filtered between 0.1 and 30 Hz. Epochs were generated at a sampling frequency of 101.73 Hz (i.e., twenty-fold decimation against raw data). For RSA analyses, epochs were generated time-locked on stimulus onset with a duration of 0.8 s (i.e., from 0.2 s before the event to 0.6 s after the event) and were baselined using the 0.2 s time window preceding the visual stimulus onset. For decoding analyses and their temporal generalization, epochs were generated from 4 s before and 4 s after stimulus onset, and were not baselined in order to avoid eliminating any anticipatory predictive activity effects. For one participant with unusable practice session MEG data, the first learning block was used in its place when necessary.

### Source reconstruction

Participants’ MEG data were co-registered to their meshed structural MRIs obtained from Freesurfer^65,66^ through identification of the same anatomical landmarks (left and right pre-auricular points, and nasion). For participants (*n* = 3) who did not undergo MRI acquisition, MEG data were morphed to a standard template brain from the FreeSurfer package. The Schaefer atlas^67^ was used to parcellate the brain into 200 cortical regions. These regions were clustered into 7 different networks: the visual, sensorimotor, dorsal attention, salience, limbic, central executive, and default mode networks. Additionally, we included 3 subcortical regions using the Desikan-Killiany parcellation^68^: the hippocampus, the thalamus, and the cerebellum. These 10 networks and regions served as our regions of interest (ROIs). For RSA analyses, noise covariance was computed with the 200 ms preceding stimulus onset, while data covariance was computed on post-stimulus onset data of each epoch. For temporal generalization and decoding analyses, noise covariance was computed on random trials only in the 200 ms preceding stimulus onset, while data covariance was computed on whole respective epochs. To estimate the time series in each ROI, the linearly-constrained minimum variance (LCMV) beamformer was computed on single-trial data. This particular spatial filter was chosen due to its ability to resolve deeper sources and suppress external noise on time series data^31^. Vector source estimates were used to keep the information of all (x, y, z) directions of the dipoles. We then performed Singular Value Decomposition in a time-resolved manner to only keep the most informative direction at each time point of the source estimates.

### Representational similarity analysis

An Ordinary Least Squares (OLS) linear regression was applied to each sensor/vertex and to each time point to compute the model coefficients that capture the neural response patterns associated with each stimulus^69^. The model residuals from the OLS were used to estimate the covariance matrix using Ledoit-Wolf shrinkage for more stable covariance estimates^70^. The cross-validated Mahalanobis distance was selected as the dissimilarity metric due to its sensitivity in multivariate statistical testing for RSA in neuroimaging data^34,71,72^:

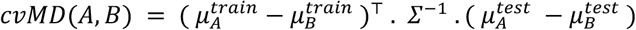

where *μ* are class means estimated separately in training and testing sets, *Σ* is the noise covariance estimated from training residuals using Ledoit-Wolf shrinkage. Cross validation was done in a leave-one-block-out manner within each run comprising 5 blocks each. This resulted in time-resolved representational dissimilarity matrices (RDMs) for each participant and each block. For each participant, the distance between elements of pattern and random pairs was extracted from the RDMs. Then the similarity index was computed using the following formula:

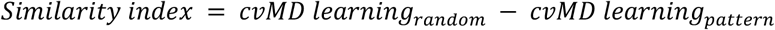

This index reflects the similarity between pair elements within pattern trials relative to the similarity between pair elements within random trials. Only pairs that appeared according to the sequence in pattern trials were considered as pairs in the random trials for the analysis (i.e., four possible pairs out of six). A positive value for the similarity index means that the similarity between elements within pattern pairs is higher than the similarity among elements within random pairs. Spearman’s ρ correlation was also computed in a time-resolved manner between the similarity and learning indices of each participant. The values were participant-centered to isolate intra-individual associations. The resulting ρ for each participant were tested against zero using a one-sample *t* test. This pipeline was applied both in sensor (whole brain) and source space (ROIs).

### Decoding

Except for correlation analyses, only learning block trials were considered in order to prevent dampening of found effects by the practice blocks. Multivariate pattern analysis (MVPA) decoding aimed at predicting the value of a specific variable *y* (i.e., the spatial arrows) from the brain signal *X*. The multiclass logistic regression was selected as the classifier. The data (*X*) were whitened using a standard scaler that *z* scored each channel at each time point across trials. Each estimator was fitted on each participant separately, across all MEG sensors/source vertices, and at a unique time sample, using the default parameters of scikit-learn. The cross-validation was performed using a leave-one-block-out (LoBo) stratified folding across all blocks, and accuracy was selected as the metric used to assess classifier’s performance. This pipeline was applied in source space only.

### Temporal generalization

A classifier (see “Decoding” section) that has been trained at a specific time sample was tested at all possible time samples using the temporal generalization method^29^ and the LoBo cross-validation strategy, resulting in two-dimensional arrays of decoding performance in accuracy (i.e., time generalization matrix) for each participant at each block. This was done for both pattern and random trials. To test if the group-mean accuracy of the decoding performance is above chance level (25%) at each time point, a non-parametric *t*-test corrected for multiple timepoint comparisons with cluster-based permutations was computed^73^. Next, the contrast was computed by subtracting the random generalization matrix from the pattern one to evidence any predictive activity prior to stimulus onset. To test if this difference in accuracy was significant (above 0), we performed a non-parametric *t*-test corrected for multiple comparisons with cluster-based permutations^73^. This pipeline was applied in sensor space only.

### Non-linear temporal modelling

One-dimensional time series such as the temporal evolution of behavioral performance, predictive activity and representational similarity over the time course of the experiment (Fig. 6 and 7, Fig S7) as well as the time course of predictive activity and representational similarity across peristimulus windows (Fig. 3-5) were investigated using generalized additive mixed models (GAMM) using the mgcv^74^ package (version 1.9.0) in R (version 4.3.2). Each variable was modelled with a GAMM including a single fixed effect: a smooth penalized regression spline of block (or time). Therefore, block (or time) was treated as a continuous predictor. Participant heterogeneity was captured by a random intercept and a random smooth for block (or time), allowing individual temporal trajectories to deviate from the population curve both by a constant offset and by a non-linear participant-specific function constrained to share a common smoothing parameter across participants. Thus, the models mgcv syntax followed this structure:

~~~
gam(value ∼ s(block, k = 10) + # fixed-effects smooth
 s(subject, bs = “re”) + # random by-participant intercept
 s(block, subject, bs = “fs”, m = 2), # random by-participant smooth for block
 method = “REML”)
~~~

Furthermore, in order to formally compare the temporal dynamics across the experiment of the three metrics of interest (Fig. 5 and 6), a fourth model was fitted to the standardized values. Each metric was scaled to have a unit standard deviation at the group level. The three metrics were grouped into a single factor variable which was allowed to interact with block via the addition of a factor smooth interaction in the model predictors. This way, each (standardized) metric is modelled by its deviation from the mean of the three, similar to a sum-to-zero contrast (aka sum contrast coding) in a linear model. A significant overall test for this interaction term indicates that at least one metric’s temporal trajectory differs from the others. Submodels including only two of the three metrics were fitted as post hoc tests to identify the differing pairs. All models used thin-plate regression splines and a basis dimension of 10 (the default values), and assumed Gaussian errors with an identity link. The adequacy of these parameters and validity of the Gaussian assumption was inspected with ad hoc diagnostics provided by mgcv. In particular, for all models, residuals were found to be normally distributed and estimated degrees-of-freedom of fixed effect smooths were well below 10. Parameters and smooth penalties were estimated by restricted maximum likelihood (REML, the default).

## Supporting information

All Supplementary figures and table

## Data Availability

Raw data will be made publicly available on a hosting platform prior to publication. Further information and requests for resources should be directed to and will be fulfilled by the correspondence authors. Source data are provided with this paper.

## Code Availability

All analyses pipelines are available at https://github.com/MEL-Eduwell-lab/asrt_analyses.

## Acknowledgements

We are grateful to Arnaud Rey, whose ideas and inspiring conversations were instrumental in shaping and realizing this project, and to Denis Schwartz for his valuable insights and expertise on source reconstruction. We thank all participants for their time and engagement. We are also grateful to the two anonymous reviewers for their thoughtful comments, which substantially improved the manuscript. We thank the MNE community for help and support. We gratefully acknowledge support from the CNRS/IN2P3 Computing Center (Lyon - France) for providing computing and data-processing resources needed for this work.

## Author Contributions Statement

**C.T**.: Conceptualization, Formal analysis, Investigation, Methodology, Visualization, Writing—original draft, Writing—review & editing. **O.A**.: Formal analysis, Investigation, Methodology, Visualization, Writing—review & editing. **T.V**.: Conceptualization, Software, Data curation, Writing—review & editing. **L.T**.: Data curation, Writing—review & editing. **A.B**.: Writing—review & editing. **M.V**.: Writing—review & editing. **D.N**.: Conceptualization, Funding acquisition, Methodology, Supervision, Project administration, Writing—review & editing. **R.Q**.: Conceptualization, Formal analysis, Funding acquisition, Investigation, Methodology, Supervision, Project administration, Writing—review & editing.

## Funding

This work was supported by a doctoral fellowship from L’école doctorale Neurosciences et Cognition (ED476-NSCO) of Université Claude Bernard Lyon 1 (to C.T.) and by the ATIP-Avenir program (to R.Q.). This work was performed within the framework of the LABEX CORTEX (ANR-11-LABX-0042) of l’Université Claude Bernard Lyon 1, within the program “Investissements d’Avenir” (decision n° 2019-ANR-LABX-02) operated by the French National Research Agency (ANR). This work was also supported by the French National Grant Agency (ANR-22-CPJ1-0042-01 and ANR-24-CE37-5807) (to D.N.); the National Brain Research Program project NAP2022-I-2/2022 (to D.N.). The data collection in Marseille was supported by LABEX Brain & Language Research Institute (BLRI) at Aix-Marseille University. This study was supported by Agence Nationale de la Recherche Grant ANR-11-INBS-0006 to France Life Imaging network and ANR-16-CONV-0002 to Institute of Language, Communication, and the Brain.

## Competing Interests Statement

The authors declare no competing interests.

## Notes

### Competing Interest Statement

The authors have declared no competing interest.

### Summary of Updates

Following peer review: - Additional elements to discuss predictive activity in the discussion - Added effect sizes on figures and additional info on statistical tests

