## Supplementary material for "Learning regularities in noise engages both neural predictive activity and representational changes": All Supplementary figures and table

### Supplementary information

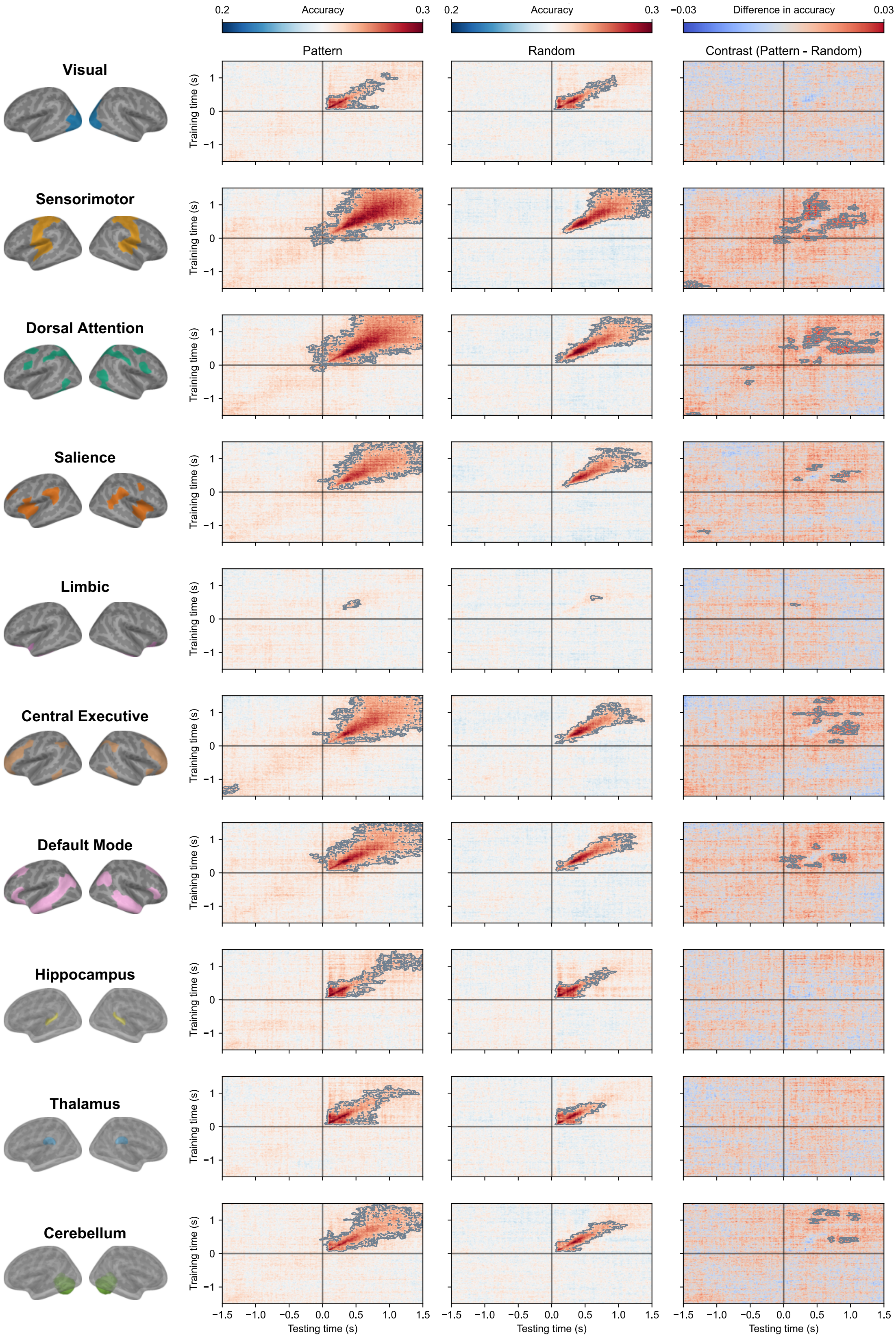


**Fig. S1 | Source space temporal generalization for pre-activation. Left** Time generalization of spatial arrows in pattern trials across all ROIs, using a logistic regression classifier and a leave-one-block-out cross-validation, averaged across learning sessions and participants. **Center** Same as **Left** but for random trials **Right** The contrast between pattern and random trials time generalization matrices. Black lines indicate the stimulus onset (0 s). Statistically significant temporal clusters (cluster-based permutation *t*-test, two-sided, $p < .01$, cluster-corrected) are contoured in grey. Data are mean across $n = 15$ participants. Source data are provided as a Source Data file.

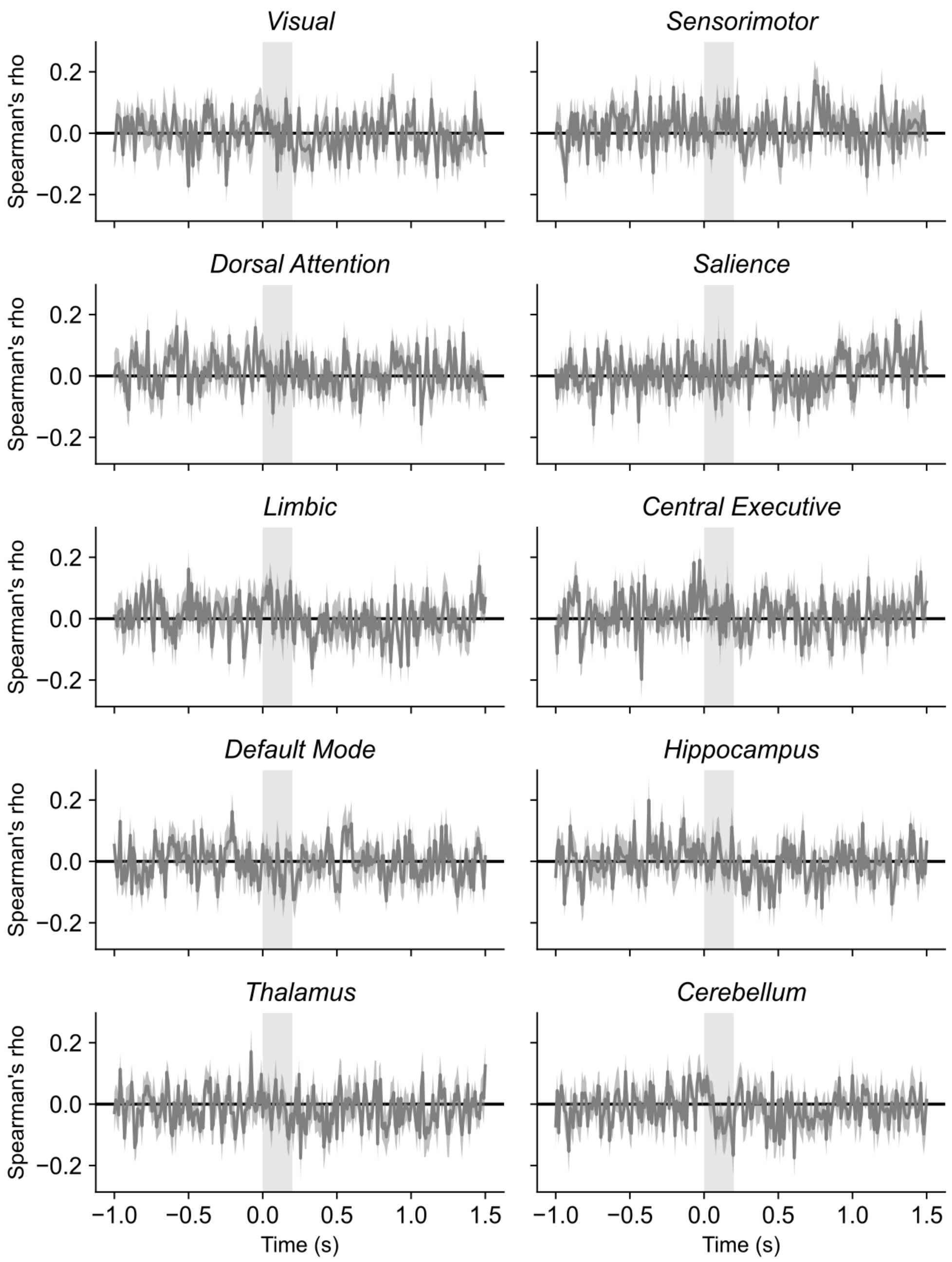


**Fig. S2 | Time-resolved spearman correlation between predictive activity and learning index across ROIs and blocks.** Grey and light grey curves represent mean Spearman's ρ value and SEM, respectively. The grey rectangle indicates stimulus duration (200 ms). Statistical significance was assessed by generalized additive mixed model (GAMM)-estimated mean differing from 0 (two-sided, $p < .05$) and overall GAMM significance against a null hypothesis of a constant zero temporal trend. No significant clusters were found in any of the ROIs (GAMM, all $p >$ .05, Holm-corrected for multiple network comparisons). Data are mean ± SEM across $n = 15$ participants. Source data are provided as a Source Data file.

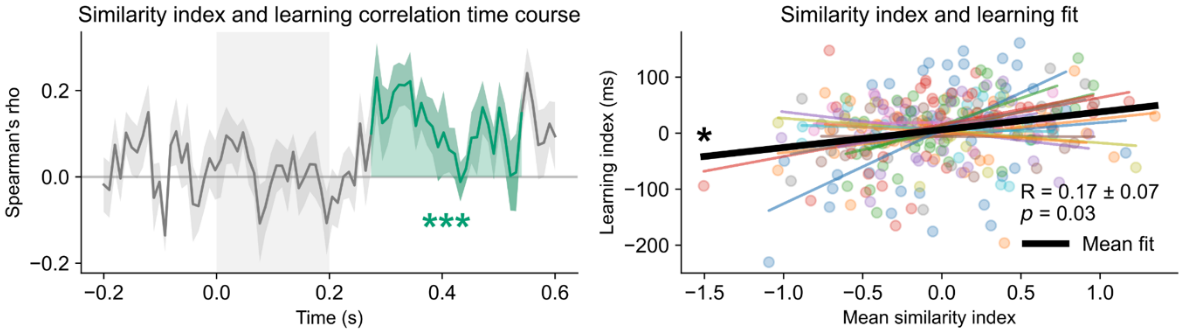


**Fig S3** | **Similarity index and learning correlation without practice blocks at sensor level. Left** Time course of Spearman's ρ correlation between the similarity and learning indices across blocks. Solid lines and shaded areas indicate the mean and the SEM, respectively. color-filled areas indicate timepoints at which GAMM-estimated mean differs significantly from zero (two-sided, $p < .05$), and asterisks indicate overall GAMM significance against a null hypothesis of a constant zero temporal trend (***$p < .001$, *$p < .05$). The similarity index positively correlates with learning during 0.27 to 0.54 s post-stimulus onset. **Right** Relationship between representational change and learning index across blocks. The mean representational change was calculated by averaging the similarity index between 0.34 and 0.48 s for each session and each participant. Each participant's fit is depicted in different colors with solid color lines indicating the mean fit across blocks. The bold black line shows the mean fit across participants. Within-participant computed Spearman ρ values are significantly different from 0 (exact $p = 0.0319$, two-sided one-sample $t$-test), confirming that increased learning is associated with greater representational change. Data are mean ± SEM across $n = 15$ participants. Source data are provided as a Source Data file.

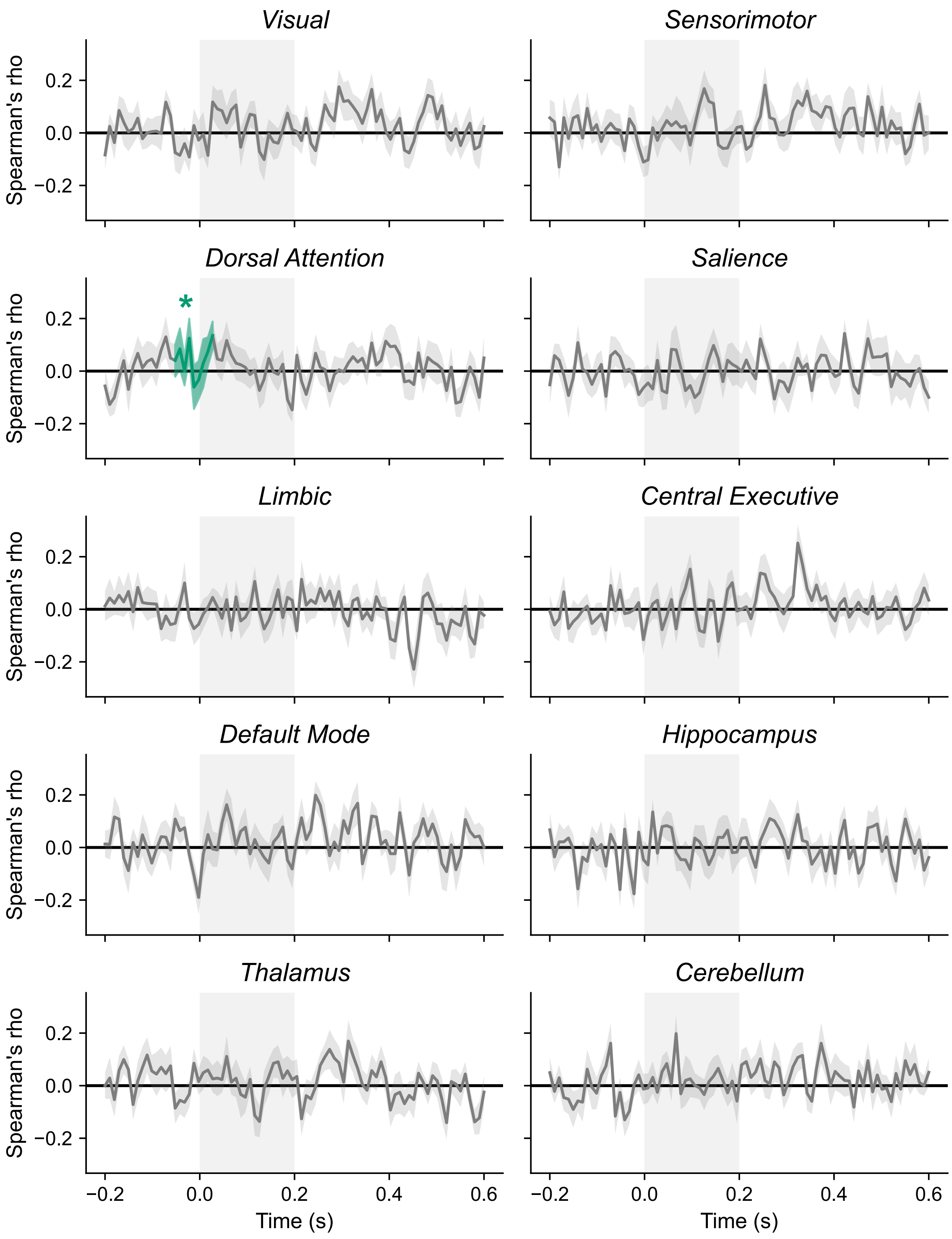


**Fig S4 | Similarity index and learning correlation without practice blocks across all ROIs.** Time course of Spearman's ρ correlation between the similarity index and the learning index across sessions. Solid lines and shaded areas indicate the mean and the SEM, respectively. The similarity index positively correlates with learning during -0.05 to 0.03 s post-stimulus onset. color-filled areas indicate timepoints at which GAMM-estimated mean differs significantly from zero (two-sided, $p < .05$), and asterisks indicate overall GAMM significance against a null hypothesis of a constant zero temporal trend (*$p < .05$, Holm-corrected across networks). No correlation is found without the practice blocks, except in the dorsal attention network before the stimulus onset which is likely a false positive due to its temporal position being inconsistent with a possible change in representational similarity. Data are mean ± SEM across $n = 15$ participants. Source data are provided as a Source Data file.

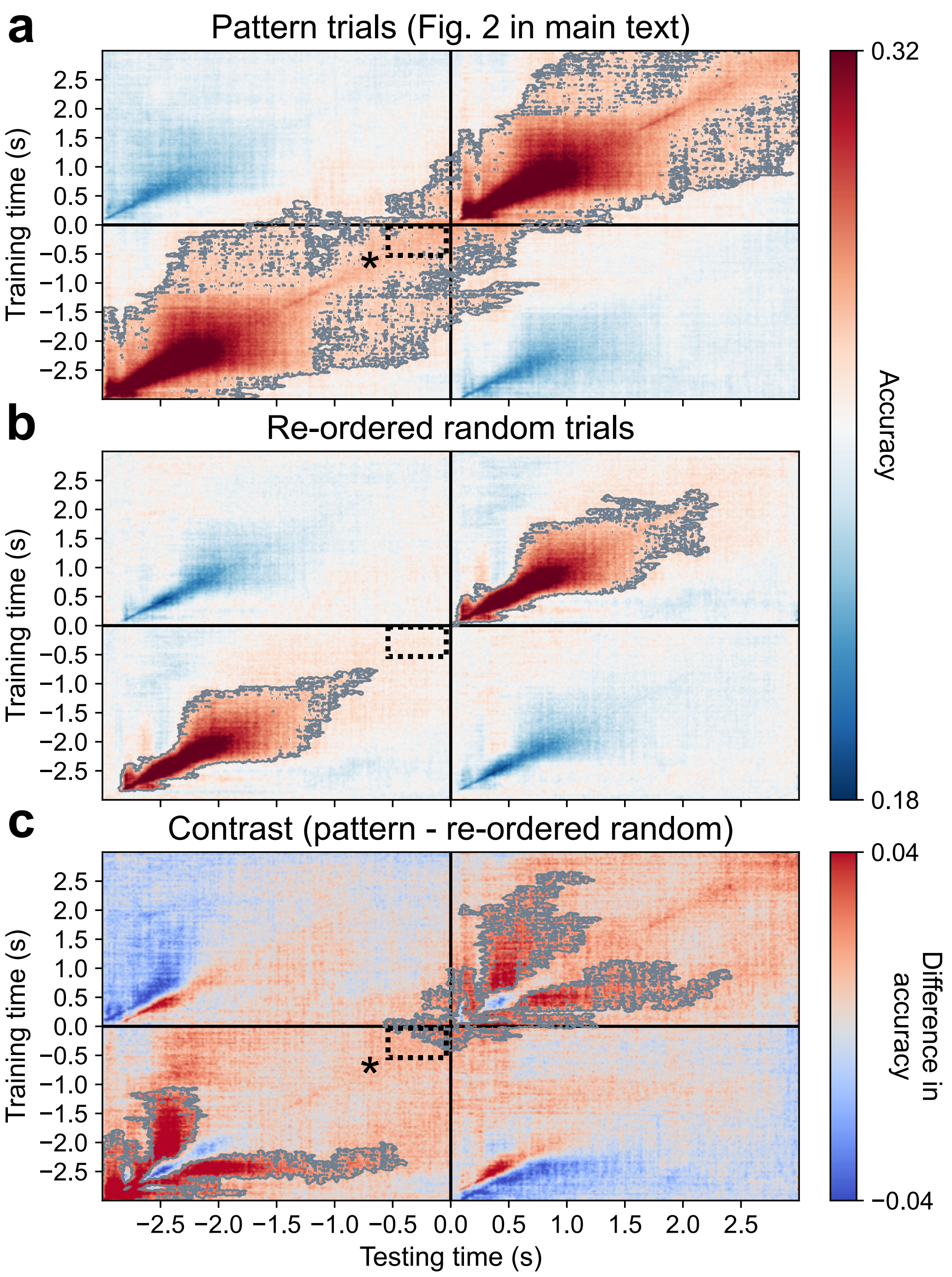


**Fig. S5 | Temporal generalization of re-ordered random trials following sequence structure. a** Time generalization of spatial arrows in pattern trials. Corresponds to Fig. 2a of the main text with an extended window of -3 to 3 s. **b** Time generalization of spatial arrows decoding in re-ordered random trials according to sequence structure. Raw data was cropped from each random trial to before the onset of the next random trial (2,900 ms later). The raw data was then re-arranged to match the pattern trials following a similar strategy than in Abdoun et al. (2026). Then we applied the same preprocessing and analysis pipelines as described in the main text. We found no neural representations during the pre-stimulus period for this reordered random condition. **c** The contrast between original pattern trials and re-ordered random trials time generalization matrices returned significant clusters in the inter-trial interval before the stimulus onset demonstrating predictive activity in pattern trials. Black lines indicate the stimulus onset timing (0 s). Statistically significant temporal clusters (cluster-based permutation *t*-test, two-sided, $p < .01$, cluster-corrected) are contoured in grey. Data are mean across $n = 15$ participants. Source data are provided as a Source Data file.

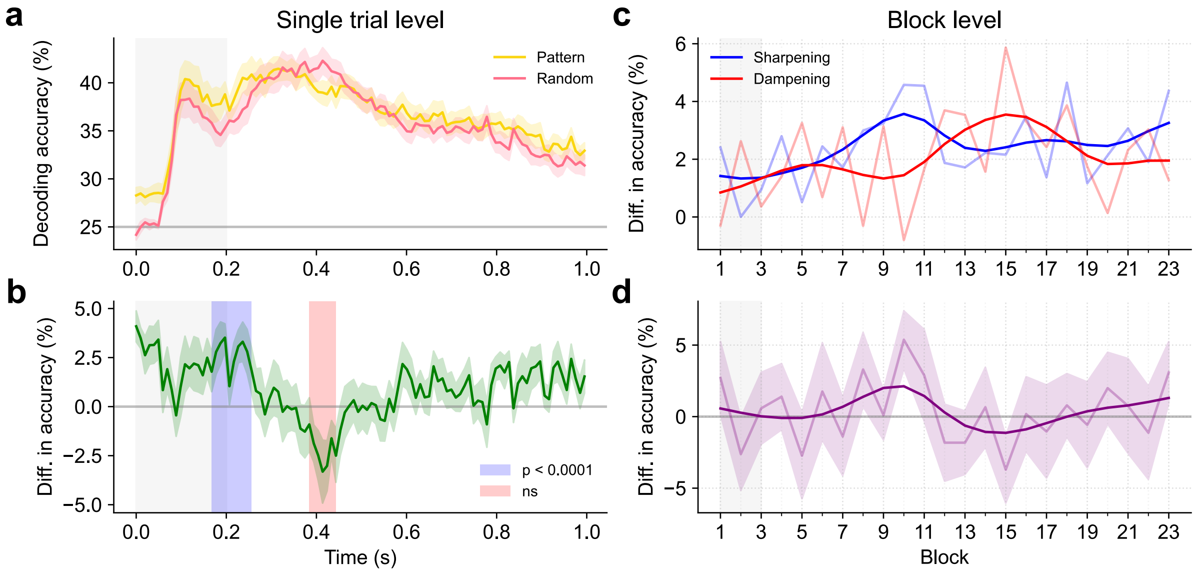


**Fig. S6 | Decoding performance in pattern and random trials. a** Decoding performance in pattern and random trials. **b** Contrast of pattern and random decoding performance. Pattern stimuli show greater decoding accuracy than random stimuli from ~0.16 to 0.26 s after stimulus onset. Random stimuli show greater decoding accuracy than pattern stimuli later from ~0.38 to 0.44 s (red area). Only the sharpening window was significant ($p < .0001$, GAMM, blue area). **c** Sharpening and dampening effects across blocks were calculated by averaging the early and late window contrast in each block. Transparent curves are mean across participants. Smoothed curves were obtained using a gaussian filter with a smoothing kernel of 1.5 blocks. **d** Contrast of sharpening - dampening across blocks. Smoothed curves were obtained using a gaussian filter with a smoothing kernel of 1.5 blocks. Sharpening shows a rapid rise peaking at block 10 then decreases and plateaus. In contrast, dampening begins its rise from block 10 onward, peaking around block 15, then decreasing. No significant clusters were found at the block-level ($p = 0.92$, GAMM). Data are mean across $n = 15$ participants. Source data are provided as a Source Data file.

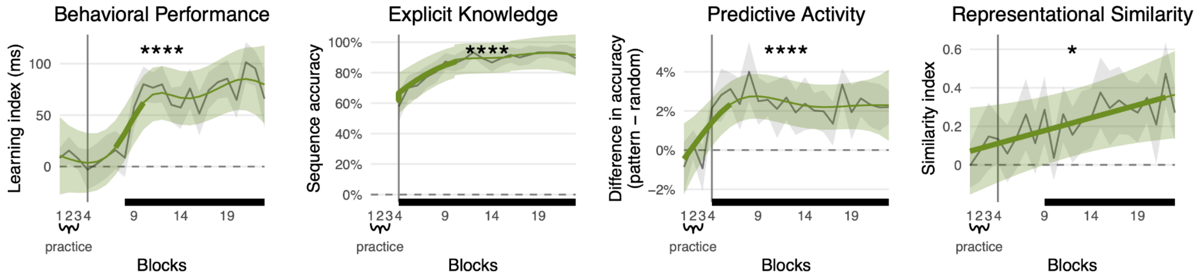


**Fig. S7 | Behavioral performance, explicit knowledge, predictive activity and representational change have different temporal dynamics across blocks.** Vertical grey lines separate practice from learning blocks. Grey lines and ribbons show the group mean ± SEM. Green lines and ribbons show the GAMM-estimated marginal means and 95% CI. Horizontal black segments mark blocks whose GAMM-estimated mean differs significantly from 0 (two-sided, $p < .05$). Thicker portions of the green smooth mark blocks with a significant temporal slope (GAMM derivative test, two-sided, $p < .05$). Asterisks indicate overall GAMM significance against a null hypothesis of a constant zero temporal trend (****$p < .0001$, *$p < .05$; Holm-corrected for metrics). Behavioral performance is significantly above 0 from block 8 ($p < .0001$, GAMM), explicit knowledge from block 4 ($p < .0001$, GAMM) predictive activity from block 4 ($p < .0001$, GAMM), and representational change from block 9 ($p = .0468$, GAMM). Data are mean ± SEM across $n = 15$ participants. Source data are provided as a Source Data file.

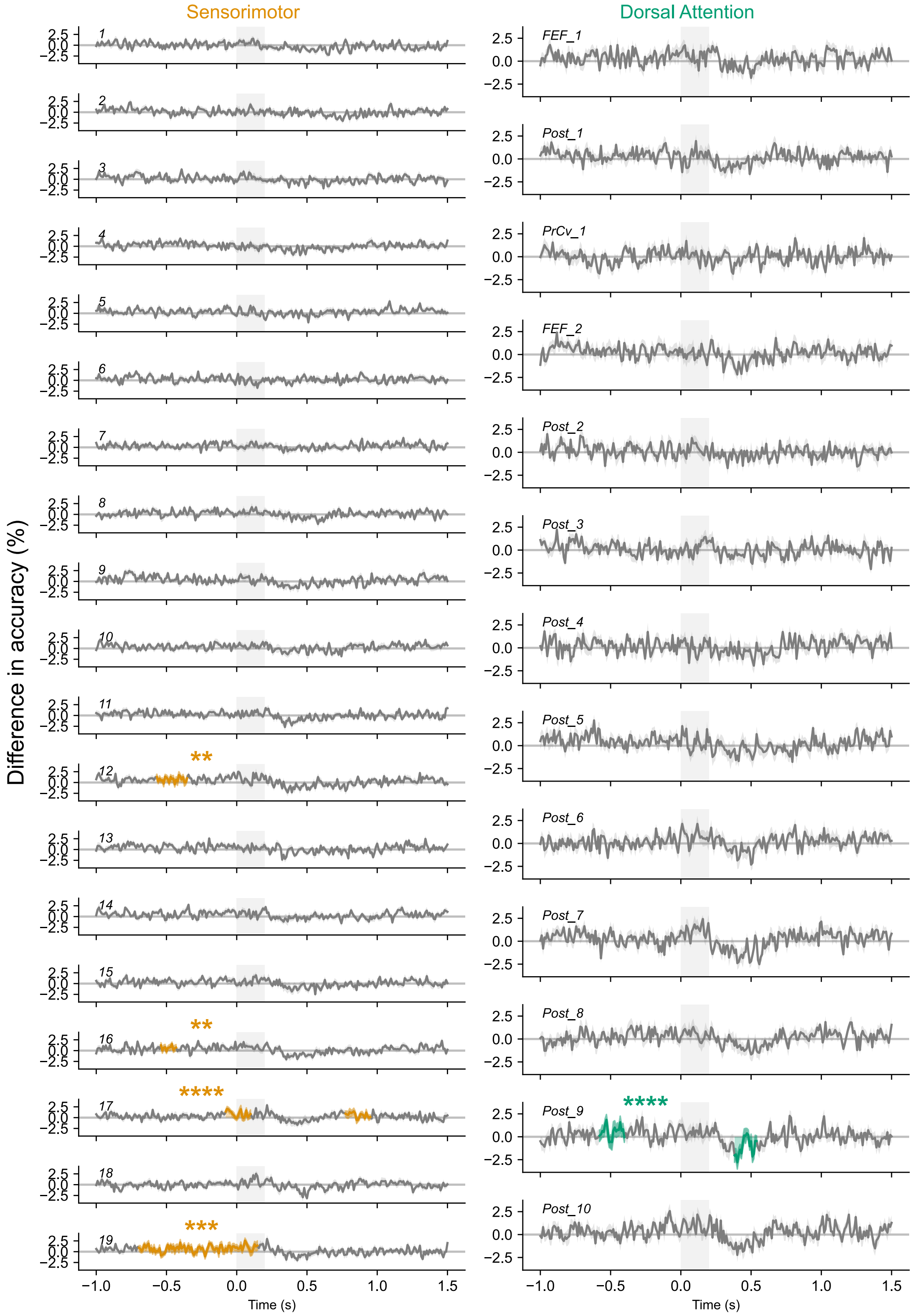


**Fig. S8 | Contrast decoding performance in parcels of Sensorimotor and Dorsal Attention networks.** Contrast between decoding performance of spatial arrows in pattern and random trials, averaged across participants and learning blocks. Left and right hemisphere parcels were merged when possible. We used a Logistic Regression classifier and Leave-one-Block-out cross-validation. The grey rectangle indicates stimulus duration (200 ms). Color-filled areas indicate timepoints at which GAMM-estimated mean differs significantly from 0 (two-sided, $p < .05$). Asterisks indicate overall GAMM significance against a null hypothesis of a constant zero temporal trend (****$p < .0001$, ***$p < .001$, **$p < .01$; Holm-corrected across networks). Data are mean ± SEM across $n = 15$ participants. Source data are provided as a Source Data file.

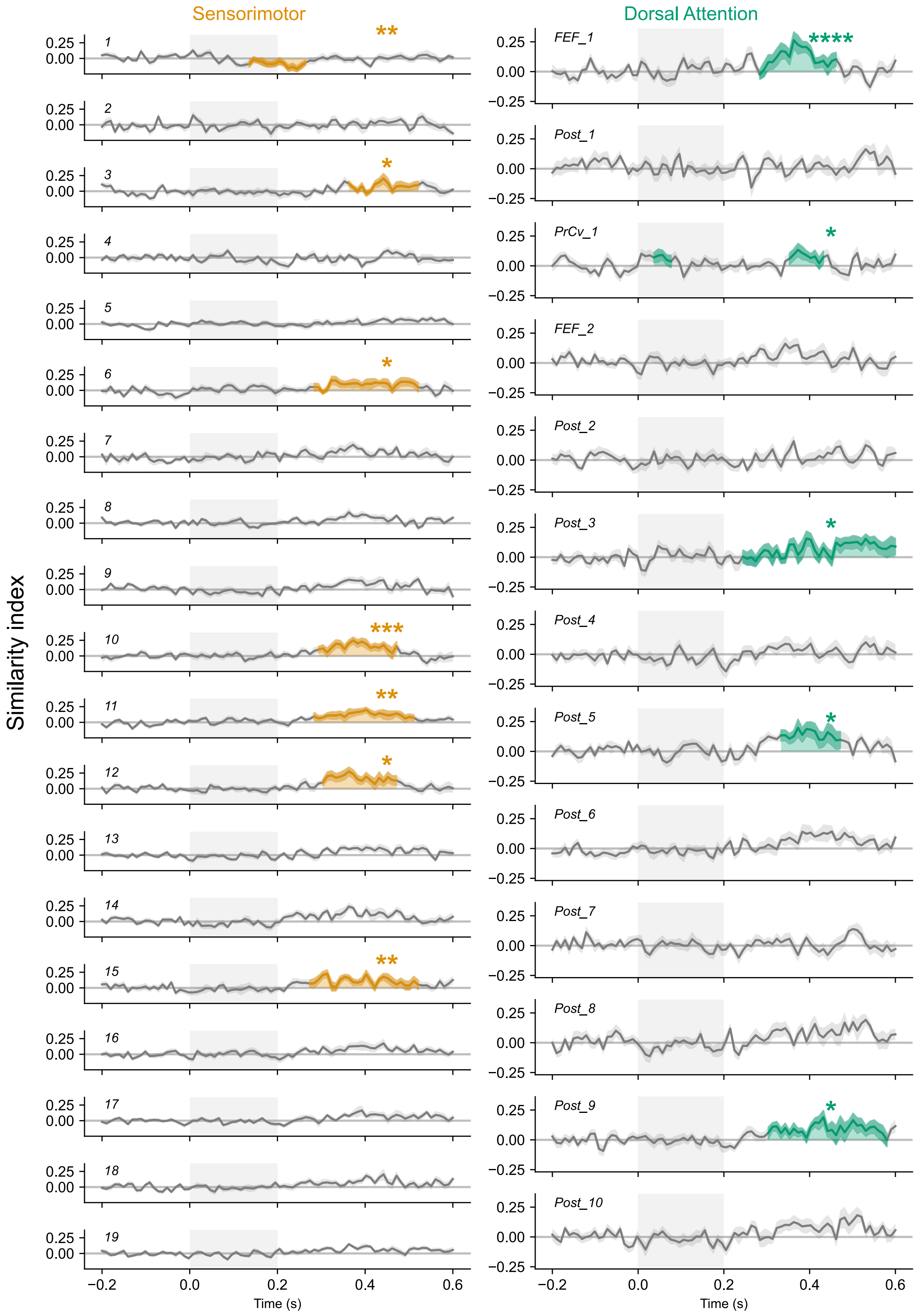


**Fig. S9 | Contrast representational similarity analysis (RSA) in parcels of Sensorimotor and Dorsal Attention networks.** Contrast between random and pattern elements in Leave-one-Block-out cross-validated Mahalanobis distance averaged across participants. The grey rectangle indicates stimulus duration (200 ms). Asterisks indicate overall GAMM significance against a null hypothesis of a constant zero temporal trend (****$p$ < .0001, ***$p$ < .001, **$p$ < .01, *$p$ < .05; Holm-corrected across networks). Data are mean ± SEM across $n$ = 15 participants. Source data are provided as a Source Data file.

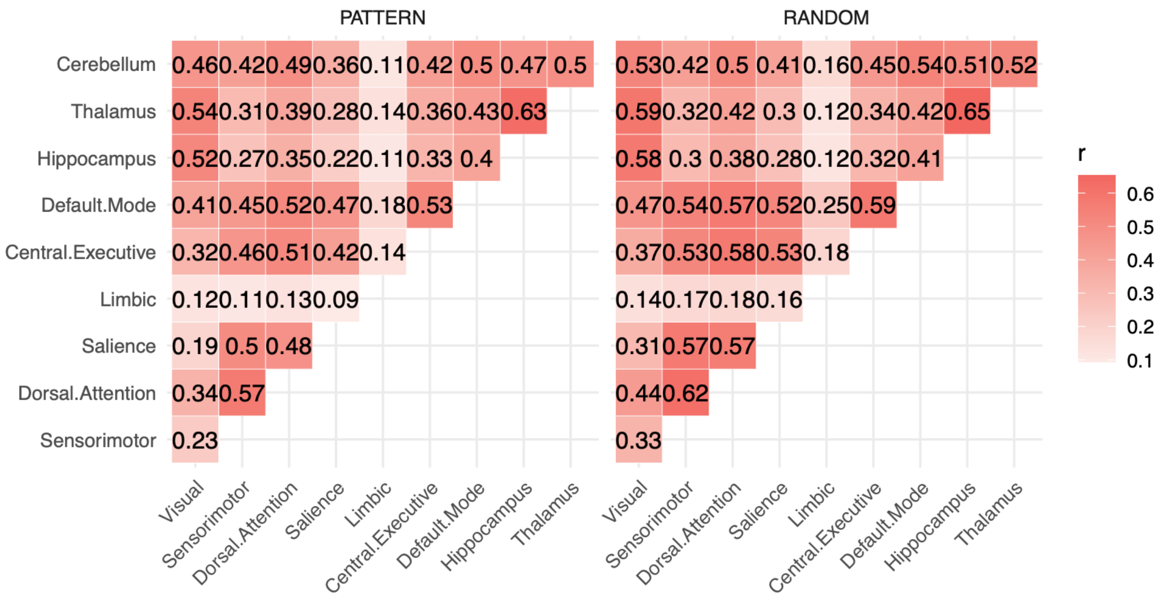


**Fig. S10 | Autocorrelation in non-linear temporal modeling of time-resolved decoding across trial type and ROIs**. (corresponds to analyses of Fig. 3a-b) Left: Pattern trials. Right: Random trials. Data are mean across *n* = 15 participants. Source data are provided as a Source Data file.

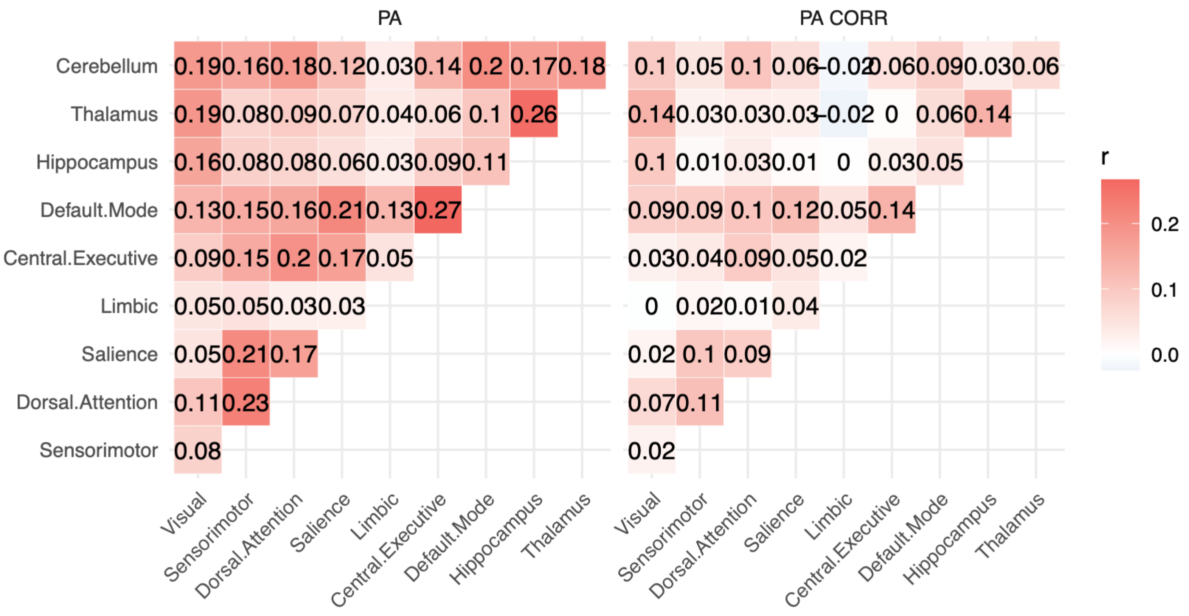


**Fig. S11 | Autocorrelation in non-linear temporal modeling time-resolved predictive activity across ROIs**. (corresponds to analyses of Fig. 3c and Fig. S3) Left: Predictive activity (PA). Right: Predictive activity correlation with the learning index (PA CORR). Data are mean across *n* = 15 participants. Source data are provided as a Source Data file.

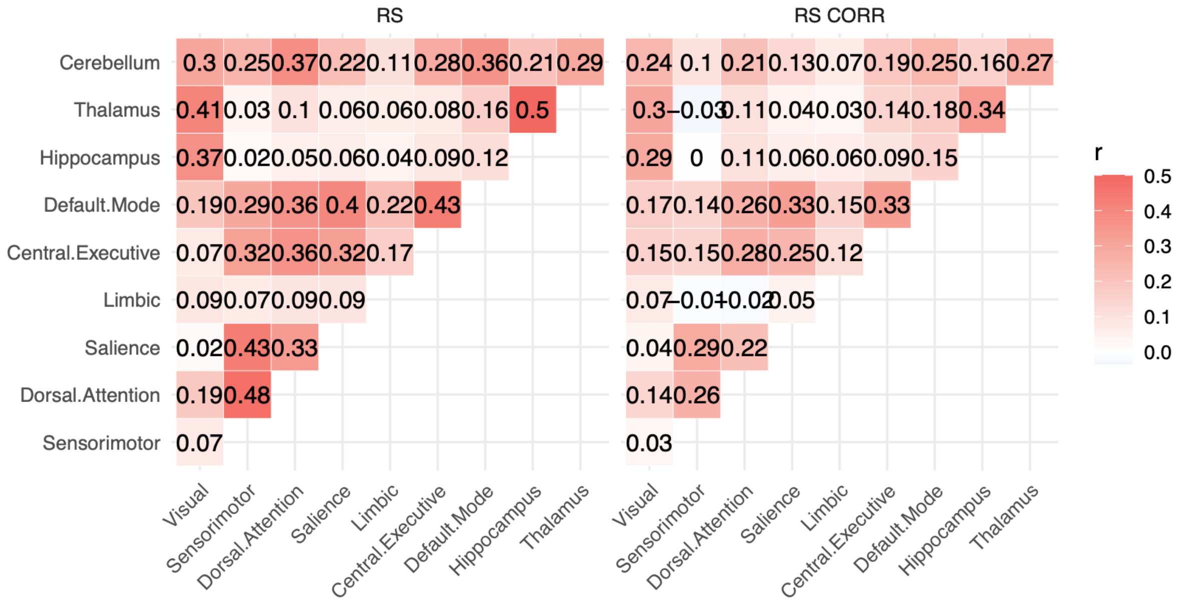


**Fig. S12 | Autocorrelation in non-linear temporal modeling time-resolved representational similarity analysis across ROIs** (corresponds to analyses of Fig. 5). Left: Representational change (RS). Right: Representational change correlation with the learning index (RS CORR). Data are mean across *n* = 15 participants. Source data are provided as a Source Data file.

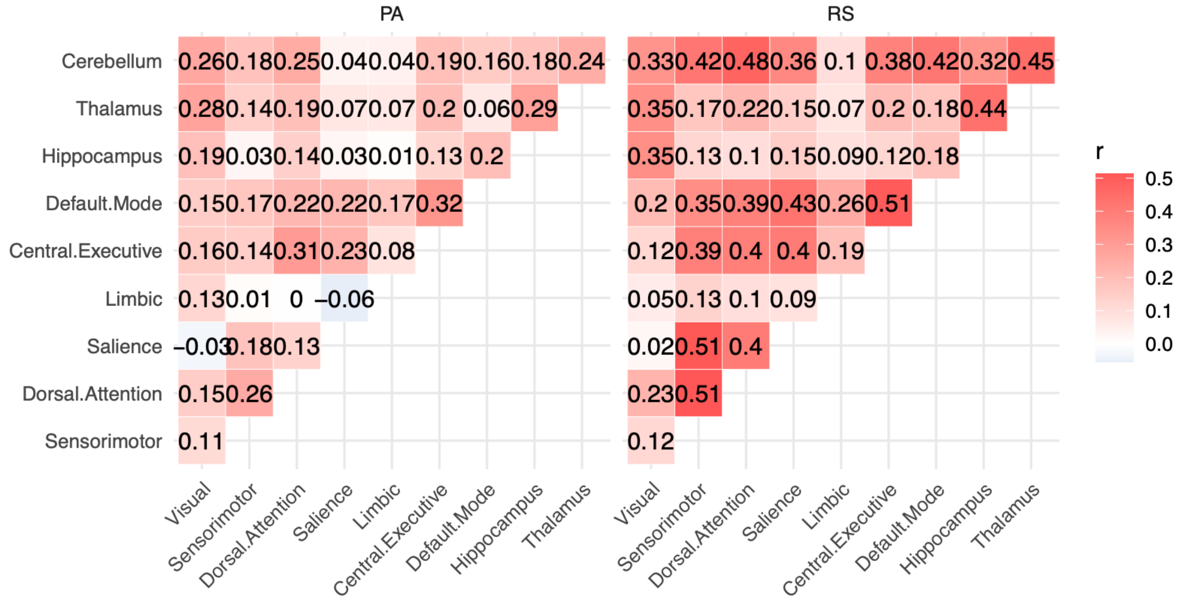


**Fig. S13 | Autocorrelation in non-linear temporal modeling of block effects across ROIs**. (corresponds to analyses of Fig. 7) Left: Predictive activity (PA). Right: Representational change (RS). Data are mean across *n* = 15 participants. Source data are provided as a Source Data file.

| | Schaefer parcel | Hemi | Destrieux label | Match (%) | MNI | | |
|---|---|---|---|---|---|---|---|
| | | | | | x | y | z |
| SENSORIMOTOR | 1 | lh | S. circular insula inferior | 35,7 | -50,8 | -9,9 | -1,8 |
| | | rh | G. temporal superior G. T. transverse | 32,4 | 50,4 | -18,8 | 5,8 |
| | 2 | lh | Lateral Fissure posterior | 28,6 | -51,7 | -27,6 | 8,1 |
| | | rh | G. temporal superior Lateral | 57,2 | 63,7 | -28,1 | 6,5 |
| | 3 | lh | Lateral Fissure posterior | 37,0 | -37,4 | -21,9 | 14,2 |
| | | rh | Lateral Fissure posterior | 55,7 | 38,1 | -16,1 | 14,5 |
| | 4 | lh | G. and S. subcentral | 74,0 | -55,5 | -6,1 | 8,7 |
| | | rh | Lateral Fissure posterior | 74,8 | 41,9 | -29,8 | 17,4 |
| | 5 | lh | G. and S. subcentral | 59,3 | -52,7 | -23,9 | 17,6 |
| | | rh | G. and S. subcentral | 76,3 | 59,4 | -2,2 | 10,3 |
| | 6 | lh | S. central | 52,4 | -54,7 | -10,5 | 29,7 |
| | | rh | G. and S. subcentral | 72,4 | 56,1 | -15,8 | 15,5 |
| | 7 | lh | S. central | 50,8 | -44,8 | -12,4 | 44,2 |
| | | rh | S. central | 53,1 | 54,3 | -8,9 | 30,8 |
| | 8 | lh | G. and S. cingulate middle posterior | 79,6 | -7,8 | -11 | 45,3 |
| | | rh | G. and S. cingulate middle posterior | 98,1 | 9,6 | -15,1 | 42,4 |
| | 9 | lh | G. postcentral | 83,6 | -46,7 | -29,7 | 53,5 |
| | | rh | G. postcentral | 62,7 | 48,6 | -25,3 | 49,4 |
| | 10 | lh | S. central | 59,3 | -39,5 | -24,1 | 54 |
| | | rh | S. central | 56,5 | 43,5 | -13,5 | 46,1 |
| | 11 | lh | S. postcentral | 55,1 | -30,5 | -45 | 58,8 |
| | | rh | G. and S. cingulate middle posterior | 66,9 | 7,4 | -10,7 | 49,6 |
| | 12 | lh | G. precentral | 58,8 | -31,9 | -21,6 | 60 |
| | | rh | S. central | 68,7 | 36,8 | -24,9 | 54,2 |
| | 13 | lh | S. postcentral | 57,6 | -26,9 | -36,4 | 63,3 |
| | | rh | S. postcentral | 48,2 | 30,6 | -42,4 | 57,5 |
| | 14 | lh | S. precentral superior part | 47,6 | -22,2 | -12,5 | 64,4 |
| | | rh | G. precentral | 54,6 | 30,6 | -21,4 | 60,1 |
| | 15 | lh | G. and S. paracentral | 58,6 | -7,8 | -31 | 65,1 |
| | | rh | S. postcentral | 58,2 | 28,1 | -36,6 | 61,9 |
| | 16 | lh | S. central | 67,5 | -19,9 | -31,1 | 63,4 |
| | | rh | S. precentral superior part | 53,9 | 22,5 | -10,2 | 63,2 |
| | 17 | rh | G. and S. paracentral | 61,4 | 9,6 | -40,8 | 67 |
| | 18 | rh | G. and S. paracentral | 52,8 | 6,4 | -25 | 66,7 |
| | 19 | rh | S. central | 63,7 | 19,2 | -30,7 | 64,9 |
| DORSAL ATTENTION | Post 1 | lh | G. occipital temporal lateral fusiform | 40,5 | -42,2 | -47 | -21,4 |
| | | rh | S. occipital temporal lateral | 41,5 | 47,4 | -56,8 | -13,7 |
| | Post 2 | lh | G. temporal middle | 46,9 | -55,4 | -60,7 | -3,9 |
| | | rh | S. temporal superior | 47,2 | 47,6 | -62,5 | 8,3 |
| | Post 3 | lh | S. intraparietal and P. transverse | 83,1 | -25,6 | -67,8 | 34 |
| | | rh | S. postcentral | 43,5 | 56,3 | -21,8 | 35,8 |
| | Post 4 | lh | S. postcentral | 67,4 | -50,9 | -28 | 38 |
| | | rh | G. parietal inferior supramarginal | 31,4 | 42,4 | -40,9 | 41,2 |
| | Post 5 | lh | S. postcentral | 99,9 | -36,6 | -36,6 | 42,6 |
| | | rh | S. postcentral | 84,1 | 37,5 | -34,8 | 42,8 |
| | Post 6 | lh | S. intraparietal and P. transverse | 74,3 | -33,6 | -49 | 43,4 |
| | | rh | G. parietal superior | 58,1 | 13,6 | -72,3 | 50,8 |
| | Post 7 | lh | G. parietal superior | 77,0 | -16,8 | -71 | 50,5 |
| | | rh | S. intraparietal and P. transverse | 84,2 | 33 | -50 | 46,7 |
| | Post 8 | lh | G. parietal superior | 61,1 | -27,8 | -59 | 55,1 |
| | | rh | S. intraparietal and P. transverse | 60,6 | 24,4 | -60,3 | 53,8 |
| | Post 9 | lh | G. precuneus | 72,9 | -7,6 | -57,8 | 56,8 |
| | | rh | G. precuneus | 73,2 | 7 | -56,1 | 59,9 |
| | Post 10 | lh | G. parietal superior | 66,8 | -18,4 | -50,3 | 66,3 |
| | | rh | G. parietal superior | 77,4 | 20 | -49,1 | 65,9 |
| | FEF 1 | lh | S. precentral superior part | 67,8 | -32,3 | -4,9 | 49,7 |
| | | rh | S. precentral superior part | 67,4 | 34 | -4,8 | 50,6 |
| | FEF 2 | lh | S. frontal superior | 64,7 | -23,1 | 5,4 | 57 |
| | | rh | S. frontal superior | 74,0 | 25,5 | 6,2 | 53,9 |
| | PrCv 1 | lh | S. precentral inferior part | 63,1 | -46,3 | 5,3 | 27,1 |
| | | rh | S. precentral inferior part | 54,5 | 49,2 | 9,3 | 21,2 |

**Table S1 | Schaefer parcellation to Destrieux labels match of Sensorimotor and Dorsal Attention networks.** Orange (sensorimotor) and green (dorsal attention) highlighted rows correspond to significant parcels in decoding (Fig. S8) or representational similarity analyses (Fig. S9). Data are mean across *n* = 15 participants. Source data are provided as a Source Data file.
